# Depletion of BBSome Subunits Alters Receptor Endocytosis and Promotes EMT via TGF-β Signalling

**DOI:** 10.1101/2024.10.23.619933

**Authors:** Carlos Solarat, Barra-Carneiro Jose, Vila-Almuíña Sara, Alvela Cristian, Barbeito Pablo, Soloshenko Margarita, Schrøder Holm Maria, Gammoh Noor, Christensen Søren Tvorup, Valverde Diana

## Abstract

**Background:** Ciliopathies are genetic disorders caused by defects in the structure or function of the cilia and their related structures. Bardet-Biedl syndrome (BBS) is a complex ciliopathy with varied symptoms, probably due to altered membrane receptor signalling pathways.

**Methods:** We investigated the role of *BBS1* and *BBS4* gene deficiencies in retinal epithelial cells using surface membrane proteomics, co-immunoprecipitation, AlphaFold structural modelling, endocytic trafficking colocalization assays, autophagy flux analysis, and epithelial-to-mesenchymal transition (EMT) functional assays including migration and wound healing.

**Results:** Deficiencies in these BBSome components led to shorter cilia and disrupted receptor endocytosis, with accumulation of TGF-β-pathway and EMT-associated proteins on the plasma membrane of *BBS1* KO cells. Co-immunoprecipitation and AlphaFold structural modelling indicated that neither BBS1 nor BBS4 forms stable physical complexes with core endocytic components (EEA1, RAB11, RAB7, LAMP2), suggesting an indirect mechanism. *BBS1* KO cells showed sustained TGFBR1 recycling, low basal autophagic flux that increases upon TGF-β stimulation, and the most pronounced mesenchymal phenotype. *BBS4* KO cells showed reduced receptor recycling, elevated basal autophagic flux that declines upon stimulation, and the strongest canonical TGF-β/SMAD activation with reduced non-canonical ERK1/2 signalling.

**Conclusions:** Increased EMT markers and enhanced migration in *BBS1* KO cells highlight the role of BBSome-dependent receptor trafficking as a potential therapeutic target for retinal degeneration in BBS. The divergent autophagic and signalling phenotypes between *BBS1* and *BBS4* KO reveal subunit-specific contributions to TGF-β pathway dysregulation.

## Plain English summary

Bardet-Biedl syndrome (BBS) is a rare inherited disease that affects multiple organs, most notably causing progressive blindness through degeneration of the light-sensitive cells at the back of the eye. It is caused by defects in the proteins that build the BBSome, a complex that works at the base of tiny hair-like structures called cilia, which are found on the surface of almost every cell in the human body. We studied what happens in eye cells when two key BBSome genes, *BBS1* and *BBS4*, are switched off. We found that these cells accumulate a receptor protein called TGFBR1 at their surface because it cannot be properly internalised and recycled. This leads to excessive activation of a signalling pathway called TGF-β, which in turn triggers a process called epithelial-mesenchymal transition (EMT), where normally organised eye cells start to behave like disorganised, migratory cells. Interestingly, the two genes caused different patterns of disruption: loss of BBS1 produced the most dramatic cell shape changes and increased migration, while loss of BBS4 produced stronger activation of one branch of the TGF-β pathway but a milder overall change in cell identity. We also showed that the BBSome does not physically attach to the proteins that normally sort and route receptors inside the cell, suggesting it controls trafficking indirectly. These findings suggest that blocking the TGF-β pathway might help slow retinal degeneration in people with BBS and highlight the importance of understanding how receptor recycling goes wrong in ciliopathies.

## Background

Ciliopathies are a group of genetic disorders characterised by abnormalities in the structure or function of cilia and their associated centrosomes or basal bodies. These disorders often arise from defects in the primary cilium, a specialised, antenna-like organelle that extends from the surface of most vertebrate cells (1). The primary cilium plays a crucial role in detecting and translating external environmental signals into intracellular responses, regulating a wide range of developmental signalling pathways, such as Hedgehog (Shh), Wnt, and TGF-β/BMP (2,3). These pathways are essential for proper cellular communication, tissue development, and organogenesis. When the function of the primary cilium is disrupted, it can lead to a variety of clinical manifestations, which include most tissues and organs of the body, highlighting the importance of cilia in maintaining normal cellular and developmental processes (4,5).

Bardet-Biedl syndrome (BBS) is a multisystemic ciliopathy marked by retinal degeneration, obesity, polydactyly, intellectual disability, cryptorchidism, and renal abnormalities, with retinal disease being the most penetrant feature (6,7). BBS is associated with mutations in genes encoding proteins predominantly localised to the primary cilium, centrosome, basal body, and transition zone. Notably, eight of these genes (*BBS1*, *BBS2*, *BBS4*, *BBS5*, *BBS7*, *BBS8*, *BBS9*, and *BBS18*) encode components of the BBSome, an octameric protein complex, which functions as a cargo adaptor for the retrograde intraflagellar transport (IFT) machinery to promote exit of ubiquitinated receptors from the cilium (5,8). Consequently, mutations in BBS genes cause major changes in the ciliary membrane proteome, to some extent associated with defective Shh and G-protein-coupled receptor (GPCR) signalling (2,5). Specifically, previous research from our laboratory has demonstrated the involvement of the transforming growth factor β (TGF-β) pathway in ciliopathy development (9).

Previous studies have established a functional connection between endosomal recycling and the primary cilium. In many cell types, including retinal pigment epithelium (RPE) cells, the base of the primary cilium is situated within a specialized invagination of the plasma membrane known as the ciliary pocket (CiPo). This region is a hub for clathrin-mediated endocytosis (CME), facilitating the formation of early endosomes and driving endocytic recycling (10–12). The close association between the CiPo and these endocytic processes highlights the crucial role of the ciliary pocket in regulating the trafficking of signalling molecules and receptors, thereby influencing ciliary function and downstream cellular signalling pathways such as in Shh (13) and TGF-β/BMP signalling (12,14). Additionally, functional connections between the BBSome and the recycling endosome factor RAB11 have been identified. ARL13B, another ciliary protein, is critical for endocytic recycling (15), underscoring the role of ciliary proteins in the recycling and transport of endosomal cargo. Deficiencies in receptor internalisation, such as Notch, leptin, and insulin receptors, have been documented in cell lines with BBSome dysfunction (16–19).

The RPE is a layer of specialised, polarised cells between the choriocapillaris and the neural retina. This layer of cells is crucial for maintaining photoreceptor health by forming a blood-retina barrier, disposing of damaged photoreceptor segments, maintaining the retinoid cycle, and protecting against light-induced and oxidative stress (20). Disruptions in these processes, particularly in protein recycling and maintenance of RPE cell polarity, can lead to dysfunction. Among these processes, autophagy and epithelial-mesenchymal transition (EMT) are particularly noteworthy. EMT in RPE cells can result in impaired tight junctions, accumulation of misfolded proteins, and dysregulation of key pathways, such as TGF-β, Wnt, and extracellular vesicle trafficking, which are also implicated in ciliopathies (21).

This study aims to investigate the role of plasma membrane alterations in receptor endocytosis pathways that may contribute to the pathological phenotype observed in the retinas of BBS patients.

## Methods

### Cell culture and treatment

Human retinal pigment epithelial cells (hTERT RPE1, ATCC CRL-4000, female) and human embryonic kidney cells (293T, ATCC CRL-3216, female) were cultured in Dulbecco’s Modified Eagle Medium F-12 nutrient mix (DMEM/F-12) supplemented with 10 % fetal bovine serum (FBS) and 1 % penicillin/streptomycin (P/S), when appropriate. Hygromycin B (0.01 mg/ml) was added for RPE1 cells. Cultures were maintained at 37 °C with 5 % CO_2_ for up to one month. Cells were 24 h serum-depleted before stimulation with TGF-β (2 ng/ml) or EGF (20 ng/ml) to enhance ciliogenesis. Bafilomycin A1 (20 nM) was used as an autophagy inhibitor.

### CRISPR/Cas9 assay and editing validation

RPE1 cells were edited using a lentivirus system according to the protocol described by other labs (22,23), with some modifications. After lentiviral transduction, cells were selected with puromycin (50 μg/ml) for 5 days.The sgRNAs used to generate the BBS1 and BBS4 knockout RPE1 cell lines and the primers used to verify CRISPR-Cas9 editing outcomes are listed in Supplementary Tables 1 and 2, respectively.

Surviving cells were expanded and isolated to obtain individual clones. The isolation was performed using Fluorescence-Activated Cell Sorting (FACS) with Hoechst (BD Bioscience) in the flow cytometer FACS ARIA III (BD Bioscience, San José, United States). Inhibition of gene expression was validated by qPCR in QuantStudio3 (Thermo Fisher Scientific). Primer information can be checked in Supplementary Tables 1 and 2. Those clones with gene expression less than or equal to 11% were tested by Western blot. Finally, those that did not present a band in the size corresponding to the BBS1 and BBS4 proteins were selected.

### RNA extraction and RT-PCR

RPE1 cells were seeded in 6-well plates at a concentration of 1.5 × 10^5^ by triplicate in DMEM/F-12 1 % P/S and were incubated for 48 h for serum depletion. Then, cells were stimulated with rhTGF-β1 (2 ng/ml) for 30 mins. DMEM was removed, and wells were washed twice with PBS. Cells were scraped in PBS and collected in 1.5 mL tubes. For RNA extraction, the NYZ total RNA isolation kit (NYZtech) was used following the manufactureŕs protocol. This process was repeated with cells without stimulation. After RNA elution, the sample concentrations were measured with Nanodrop (Thermo Fisher Scientific). Next, 100 ng of RNA was retrotranscribed to cDNA using the NZY First-Strand cDNA Synthesis kit (NYZtech) following the manufacturer’s protocol.

### qPCR expression analysis

qPCR was performed on samples prepared in 96-well plates using PowerUp SYBR Green Master Mix and QuantStudio3 (Thermo Fisher Scientific). Gene expression was normalised using the ΔΔCT method with *ALAS1*, *YWHAZ,* or *B2M* as housekeeping genes. Each sample was analysed in technical triplicate for two biological replicates (clones 1 and 2). Primers can be found in Supplementary Table 3.

### Fluorescence imaging

Cells were grown on glass coverslips in a 6-well plate. 48 h later, cells were treated as indicated. Coverslips were then fixed with 4 % ice-cold paraformaldehyde (PFA) in 20 mM HEPES, pH 7.5, for 10 min. Cells were permeabilised in 0.1 % Triton X-100 in PBS for 5 min at RT. Slides were then incubated in primary antibodies in blocking buffer (1 % BSA in PBS) at 37 °C for 2–3 h or overnight at 4 °C, followed by incubation with Alexa-conjugated secondary antibodies (Invitrogen) for 1 h at room temperature. Finally, nuclei were stained with 1 μg/ml DAPI (Sigma) for 5 min at RT; then, coverslips were mounted on microscope slides with Prolong Diamond Anti-fade (Invitrogen).

### Co-immunoprecipitation and structural modelling

#### Longer but more accurate version

HEK239T cells were transfected with pcDNA-Flag-HA-BBS1, pcDNA-Flag-HA-BBS4 or an empty vector using the calcium phosphate method and lysed 40–48 hr later in EBC buffer (50 mM Tris-HCl pH 7.5, 150 mM NaCl, 1% Triton-X-100) and 1 X Halt protease and Phosphatase inhibitor cocktail (Thermofisher, #78440). Protein levels in the total protein fraction were measured with the Pierce BCA Protein Assay Kit (Thermofisher), and the protein amount and concentration were equalised in all samples. HA-tagged BBS1 or BBS4 was immunoprecipitated using anti-HA antibody (Abcam, ab9110) and Dynabeads® Protein A (Thermofisher, 10002D) following manufacturer’s instructions. Beads were then washed thrice in lysis buffer without inhibitors, eluted with 2 X Laemmli buffer containing β-Mercaptoethanol (BME), and boiled for 5 min at 95 °C. The total protein fraction, supernatant fraction, and immunoprecipitated fraction were resolved by SDS–PAGE and probed by Western blot for HA, EEA1, RAB11, RAB7, and LAMP2.

For structural modelling, we used AlphaFold 3 (v3.0.1) as a primary screen of pairwise interactions between BBSome subunits and candidate GTPases, with AlphaFold 2-Multimer (v2.3.2, full databases) used as cross-validation for ambiguous pairs. For AlphaFold 3, five diffusion samples × 10 recycles were generated per pair (seed 42, NVIDIA A100 GPU); the top-ranked model of ambiguous pairs was additionally inspected visually in PyMOL. Calibration was anchored on three controls: BBS1–ARL6 (positive; PDB 4V0O), BBS1–BBS9 (positive, intra-BBSome contact), and BBS1–GFP (negative). Based on these controls, we defined empirical thresholds of AlphaFold 3 ipTM ≥ 0.65 as positive, 0.30–0.65 as ambiguous (requiring AlphaFold 2-Multimer validation), and < 0.30 as negative, with an inter-chain Predicted Aligned Error (PAE) < 5 Å required for high-confidence interfaces. AlphaFold 2- Multimer confidence was assessed using the ranking confidence score of the top-ranked model, defined as 0.8·ipTM + 0.2·pTM. When AlphaFold 3 predictions were ambiguous, pairs were treated as computationally unsupported if AlphaFold 2-Multimer ranking confidence remained low (<0.40), for example BBS2–RAB11B (0.27). These cut-offs were applied only as pragmatic screening criteria and were not interpreted as definitive evidence for either interaction or non-interaction. Low confidence therefore indicates absence of robust structural support in this modelling framework, but does not rule out transient, indirect, post-translationally regulated, membrane/context-dependent, or oligomer-state-dependent associations. Using this pipeline, we screened RAB7A and RAB11B (the GTPases probed by co-immunoprecipitation) against BBS1, BBS4, and the remaining BBSome subunits.

### Membrane biotinylation assay

RPE1 cells were seeded in 100 mm dishes in triplicate in DMEM/F-12 1 % P/S, 10 % FBS. After that, DMEM was removed, and wells were washed twice with ice-cold PBS and labelled with 0.5 mg/ml biotin (EZ-Link sulfo-NHS-SS-biotin, Thermo Fisher Scientific) for 30 min at 4 °C. Excess sulfo-NHS-SS-biotin was washed off with PBS and then quenched with 100 mM glycine for 15 min at 4 °C. To isolate biotinylated proteins, cells were lysed in RIPA buffer with proteasome, protease, and phosphatase inhibitors and then incubated overnight at 4 °C with Streptavidin Sepharose High Performance (Cytiva). To load the same amount of protein onto the beads, protein quantification was performed by Pierce™ BCA Protein Assay Kit (Thermo- Fisher Scientific) microplate assay (Bio-Rad). Beads were washed three times with RIPA and then once with RIPA supplemented with 300 mM NaCl final concentration, followed by two washes with PBS. Beads were then analysed by SDS–PAGE and Western blotting or prepared for mass spectrometer analysis.

RIPA composition: TrisHCl (pH 7.4) 50 mM, NaCl 150 mM, EDTA 5 mM, EGTA 1 mM, NP-40 alternative 1 %, Na desoxycholate 0.5 %, SDS 0.1 %.

### Protein extraction for proteomic profiling

For the plasma membrane proteomic profile, beads were resuspended in 100 μL 2 M urea, 100 mM Tris (pH 7.5), 1 mM DTT buffer, and 2 μg/mL trypsin and incubated at 37 °C 30 min in a shaker (1000 rpms). After that, the beads were spun at 8,000 rpm for 1 min at RT. The supernatant was kept in a new low-protein-binding microcentrifuge tube and incubated overnight at 37 °C. The next day, 5 μL of 10 % TFA (trifluoroacetic acid) was added to stop the reaction. Samples were aliquoted and stored at −80 °C or directly sent to the core facility for C18 purification before mass spec analysis.

### Mass spectrometry and bioinformatic analysis

The desalted and lyophilized peptides were resuspended in 0.1 % TFA and subjected to mass spectrometric analysis by reversed-phase nano–liquid chromatography-tandem mass spectrometry (LC-MS/MS). Mass spectrometry: 5 µl of the resuspended peptides were analyzed by reversed-phase nano–LC-MS/MS using a nano-Ultimate 3000 LC system and a Lumos Fusion mass spectrometer (Thermo Fisher Scientific). Flow rates were 400 nl/min. Peptides were loaded onto a self-packed analytical column (uChrom 1.6, 0.075 mm by 25 cm) using a 67-min gradient buffer A (2% acetonitrile, 0.5% acetic acid) and buffer B (80% acetonitrile, 0.5% acetic acid); 0 to 16 min: 2% buffer B, 16 to 56 min: 3 to 35% buffer B, 56 to 62 min: 99% buffer B; 62 to 67 min 2% buffer B. Full-scan spectra recording in the Orbitrap was in the range of m/z 350 to m/z 1,400 (resolution: 240,000; AGC: 7.5e5 ions). MS2 was performed in the ion trap, with an isolation window of 0.7, an AGC of 2e4, an HCD collision energy of 28, a rapid scan rate, a scan range of 145 to 1,450 m/z, a 50-ms maximum injection time, and an overall cycle time of 1 s.

After PEAKS imputation, an in-house limma pipeline developed by Wasim Aftab (available at [https://github.com/wasimaftab?tab=repositories](https://github.com/wasimaftab?tab=repositories)) was used to log-transform and impute the provided data. The imputation process utilized a downshift of 3 and a width cutoff of 0.3. Differential expression analysis between the two groups was performed using the Empirical Bayes Statistics method within this pipeline. The GO Terms analysis was performed with the Clusterprofiler package (24), and results were visualized with the GOplot package (25).

### TGF-β activation assays with SDS-PAGE, Western Blotting, and quantification

For TGF-β activation assays, RPE1 cells were plated in 6-well plates. The cells were initially incubated under two conditions: in DMEM without FBS and in DMEM supplemented with 1 % P/S and 10 % FBS. This incubation period lasted for 48 h, except for the SMAD2/SMAD3/ERK1/2 activation time-course, for which cells were serum-starved for 24 h as described above. Following this, the TGF-β pathway was activated by adding recombinant human TGF-β1 (rhTGF-β1) ligand at a final concentration of 2 ng/ml to each well. For the EMT marker panel, cells were stimulated for 0 and 30 min; for the SMAD2/SMAD3/ERK1/2 time-course, cells were stimulated for 0, 5, 10, 15, 30, and 60 min; for the autophagy time-course, cells were stimulated for 0, 30, and 60 min, with an additional 60 min condition in the presence of Bafilomycin A1 (20 nM). After the designated stimulation period, the cells were washed three times with PBS to remove excess ligands. The cells were then harvested using a scraper and collected by centrifugation at 7,400 × g for 10 min at 4 °C.

The resulting cell pellets were lysed on ice for 10 min in 200 μL of RIPA lysis buffer with inhibitors. The lysates were then centrifuged at 14,500 × g for 30 min at 4 °C, and the supernatant was carefully collected into 1.5 mL low-binding tubes. The remaining pellet was discarded. Protein concentrations were determined using the Pierce™ BCA Protein Assay Kit (Thermo Fisher Scientific), and the samples were subsequently stored at −80 °C until they were ready for further analysis.

For SDS-PAGE and Western blotting, 15 μg of total protein from each sample was prepared by mixing with 6.25 μL of Laemmli Buffer 4X (Bio-Rad), 1.25 μL BME, and an appropriate volume of distilled water to achieve a final volume of 25 μL. These samples were then denatured by boiling at 95 °C for 5 min. The denatured samples were loaded onto a Mini- PROTEAN® TGX™ Precast Gel with 10 % acrylamide/bis-acrylamide concentration (Bio-Rad) and subjected to electrophoresis at 150 V for 45-60 min to separate the proteins based on their molecular weight.

After electrophoresis, the separated proteins were transferred to a 0.2 μm PVDF membrane using the Trans-Blot Turbo transfer system (Bio-Rad) according to the mixed molecular weight protocol. The membranes were blocked in a solution of Tris-buffered saline containing 0.1 % Tween-20 (TBS-T) and 5 % milk for 90 min at room temperature to prevent non-specific binding. Following blocking, the membranes were incubated with the primary antibody overnight at 4 °C.

The next day, the membranes were washed three times with TBS and then incubated with the appropriate secondary antibody in TBS-T 5 % milk for 1 h at room temperature. After a further three washes in TBS-T, the protein bands were visualized using the Clarity Western ECL substrate (Thermo Fisher Scientific) and detected using the ChemiDoc system (Bio-Rad).

To assess multiple proteins on the same membrane, the membranes were stripped using Restore™ PLUS Western Blot Stripping Buffer (Thermo Fisher Scientific) according to the manufacturer’s instructions. This process involved incubating the membranes in a stripping buffer for 15 min, followed by three washes with 1X TBS. The membranes were then re-blocked and re-probed with different primary and secondary antibodies. This stripping and re-probing process was repeated up to three times.

### Transwell assay

RPE1 cells were cultured in T75 flasks and serum-depleted for 48 h once they reached 70-80 % confluency. Cell Culture Inserts for 24 Well Carrier Plate System with 8 µm pore were placed in a 24-well plate, and test media was added to the outer compartment of each well. Test media could be DMEM 10 % FBS, DMEM supplemented with 2 ng/ml TGF-β, DMEM supplemented with 2 ng/ml EGF, or DMEM alone. Then, cells were detached with 100 μL/cm2 TrypLE™ Express (Gibco), which was neutralized by adding the same number of serum- depleted media and gently shaking the culture vessel for 30 seconds. Cells were seeded in the inner compartment of each well with a density of 5 x 10^4^ cells/well. The 24-well plate was incubated in a humidified incubator (37 °C, 5 % CO2) for 6 h; after that, inserts were discarded. Then, migrated cells were washed with PBS and fixed with ice-cold methanol for 20 mins. The plate was left to dry for 30 mins after removing the methanol and incubated overnight at 4 °C. The next day, cells were stained with Crystal Violet (Merck) for 20 mins. After washing the cells three times with distilled water, cells were lysed with 10 % acetic acid, and the absorbance was measured in a microplate reader (BioRad) at a wavelength of 595 nm.

### Antibodies

Antibodies were used for either Western blotting (WB), Immunoprecipitation (IP) or immunofluorescence microscopy assay (IFM) as indicated in Supplementary Table 4. Secondary antibodies were used at 1:600 for the IFM protocol or at 1:3000 for WB. They are also present in the specific part of the STAR Methods table.

### Data acquisition and statistical analysis

Unless otherwise specified, each biological replicate consists of an independent experiment performed on separately-passaged cells. For KO cell lines, all quantitative data represent pooled results from three independent clones for *BBS1* KO and two independent clones for *BBS4* KO. Quantification of fluorescence or WB band intensity was achieved using Fiji-ImageJ software (26).Data visualization and statistical analysis were performed with the software GraphPad Prism 8.00. Differences among groups were compared using two-way ANOVA followed by Holm-Sidak correction, or Student’s t-test (two-tailed, unpaired) when there were only two groups. The differences were considered significant when ∗p < 0.05, ∗∗p < 0.01, ∗∗∗p < 0.001, and ∗∗∗∗p < 0.0001.

## Results

### Disruption of BBSome subunits causes ciliary shortening and decreased number of cilia

To study the role of the BBSome in endocytosis, we generated human retinal epithelium (RPE1) cell lines deficient in the *BBS1* or *BBS4* genes. *BBS1* was selected because it is the most frequently altered gene in the affected BBS population worldwide (the p.M390R mutation is present in 28% of patients). On the other hand, BBS4 is an external BBSome subunit that presumably interacts with many proteins involved in protein degradation, a crucial pathway in RPE1 cells for recycling photoreceptors. Furthermore, both subunits are critical for correctly assembling and translocating the BBSome to the ciliary base (27). To this end, we generated three knockout (KO) clones each harbouring a deletion of either *BBS1* or *BBS4* using the CRISPR-Cas9 gene editing technique (Suppl. Fig. 1A). Of the three clones produced for each KO, only clone 3 of the *BBS4* KO showed unsuccessful deletion. As a result, only clones 1 and 2 from each KO cell line were selected for further experimentation.

We conducted a series of experiments to confirm that our cell lines replicated the pathological phenotype described in the literature. First, we performed immunofluorescence microscopy (IFM) to determine the number and length of cilia. It is well-documented that animal models deficient for *BBS1* have a similar number of cilia to wild-type (WT) models, but they are significantly shorter (28,29). However, there is limited data for BBS4 deficiency.

We labelled ARL13B and acetylated tubulin, both recognized cilia markers, and evaluated the number and length of cilia present in the cells (Suppl. Fig. 1B). As expected, *BBS1*-deficient cells exhibited shorter cilia; in the *BBS4*-deficient cell line, the cilia were not only shorter compared to control cells but were also less frequent. It is known that *BBS1* deficiency produces a milder pathological phenotype compared to deficiencies in other BBSome subunits (30,31).

Next, we evaluated the different phases of the cell cycle in serum-supplemented media (Suppl. Fig. 1C). After 24 h of incubation, *BBS1* or *BBS4* KO cells show no significant differences compared with the WT.

We then performed an apoptosis assay to assess how the reduction in ciliary length might impact cell survival. (Suppl. Fig. 1C). The *BBS1* and *BBS4* KO cells had lower early apoptotic populations (3.8 % and 3.6 %, respectively) compared to WT cells (6.5 %) when grown in serum-supplemented media. However, in serum-depleted media, which promotes cilia formation, only *BBS4* KO cells showed a significant change in early apoptosis. These results suggest a mild resistance to apoptosis in the KO cells, indicating a differential response associated with ciliary formation.

In conclusion, our findings highlight that *BBS1* and *BBS4* deficiencies impact ciliary structure and cell survival, contributing to the understanding of their roles in ciliopathies.

### Loss of BBS proteins results in an accumulation of proteins related to the TGF-β pathway and EMT

After confirming that *BBS1* and *BBS4* deficiency impacts ciliary structure and cell survival, we aimed to identify the primary receptors affected, in addition to the known cases of Notch, insulin, and leptin receptors. In proliferating cells, the bulk of endosomal trafficking decreases, but CME is responsible for internalising specific receptors, such as those for TGF-β, fibroblast growth factor (FGFR), the Notch receptor, and components of the planar cell polarity (PCP) complex (Celsr1, Frizzled 6, Vangl2), as shown in fly models (32–36). The function of this selective receptor endocytosis is to ensure equal or asymmetric partitioning of these receptors between the daughter cells after mitosis.

To capture the broadest possible set of receptors affected by BBSome dysfunction, we performed a proteomic analysis focused exclusively on cell surface proteins. As the aim was to determine the largest number of receptors that could be affected by an inefficient function of the BBSome, these experiments were carried out in serum-supplemented media, i.e., under cycling conditions. We seeded three replicates of each of the three clones for *BBS1* KO, *BBS4* KO, and WT. The surface proteins were labelled with biotin and isolated via streptavidin pulldown.

The proteomic analysis revealed that *BBS1* KO cells exhibited an accumulation of proteins associated with all the pathways described above (FZD6, FGFR1, TGFBR1) and more internalisation of DPP4 (Figure 1A). Inhibition of the internalisation process of these receptors can disrupt the partitioning of them among the daughter cells and compromise tissue polarity (32,36), which seems to be the case in *BBS1* KO cells. One of the main characteristics of ciliopathies is loss of cell polarity (37–39).

**Figure 1.**
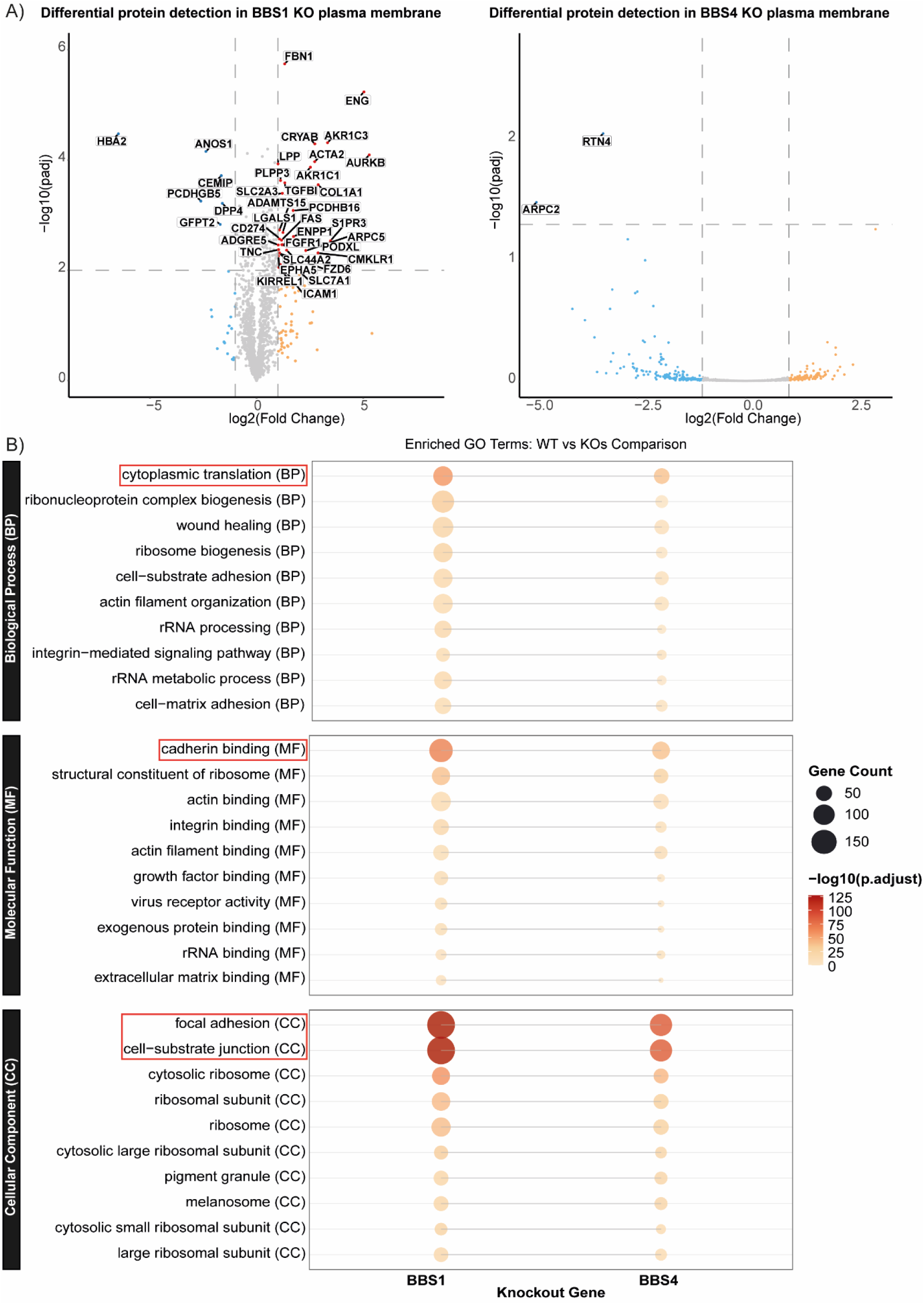
Proteomic analysis of the plasma membrane in BBS1 KO cells. (A) Volcano plot illustrating proteins overrepresented or underrepresented in the plasma membrane of BBS1 KO and BBS4 KO compared to WT cells, highlighting cell-polarity- and EMT/TGF-β-pathway-related proteins (FZD6, FGFR, TGFBR1, DPP4, ENG, CRYAB, ACTA2). A fold change threshold of ±1.5 and an adjusted p-value cutoff of 0.05 were applied for significance. (B) Gene Ontology (GO) analysis of the altered membrane protein receptors in BBS1 KO cells, showing enrichment in EMT-related processes including focal adhesion, cell-substrate junction, cadherin binding, and cytoplasmic translation.

Also, key proteins of EMT and the TGF-β pathway (Figure 1A) are accumulated, including endoglin (ENG), crystallin-α B (CRYAB), and actin alpha 2 (ACTA2), suggesting that these pathways are dysregulated in the absence of BBS1. Gene Ontology (GO) analysis of the altered membrane receptors further indicated disruptions in EMT-associated processes, including focal adhesion, cell-substrate junction, cadherin binding, and cytoplasmic translation (Figure 1 B).

To verify that the enrichment of proteins related to the TGF-β pathway was a result of the pathological phenotype associated with the KO lines, rather than transcriptional overexpression, we performed qPCR to measure the expression levels of *TGFBR1* and *IFT88*, the latter encoding a protein of the IFT-B complex essential for ciliary assembly and maintenance (40). Neither gene was overexpressed in any of the KO cells compared to the WT (Suppl. Fig. 2), indicating that the accumulation of proteins in the membranes of the KO cell lines reflects altered trafficking rather than increased transcription. Neither of these two proteins is detected in the total differentially expressed proteome when comparing KO and WT cells (data not shown).

### BBS1 and BBS4 do not form stable complexes with the endocytic trafficking machinery

Having identified that components of the TGF-β and EMT pathways accumulate at the plasma membrane of *BBS1* KO cells, we next asked whether BBS1 or BBS4 interact directly with the endocytic trafficking machinery that operates at the CiPo, as a possible explanation for this accumulation. To this end, we performed co-immunoprecipitation (co-IP) assays in HEK293T cells transfected with pcDNA-Flag-HA-BBS1 or pcDNA-Flag-HA-BBS4 plasmids, probing the total protein fraction, supernatant, and immunoprecipitated fraction with antibodies against the endosomal trafficking markers EEA1 (early endosomes), RAB11 (recycling endosomes), RAB7 (late endosomes), and LAMP2 (lysosomes), alongside an anti-HA antibody as positive control (Figure 2). A schematic of the BBS proteins and the RAB7A, RAB11B, and control ARL6, highlighting the protein domains used to guide this analysis, is shown in Figure 2A.

**Figure 2.**
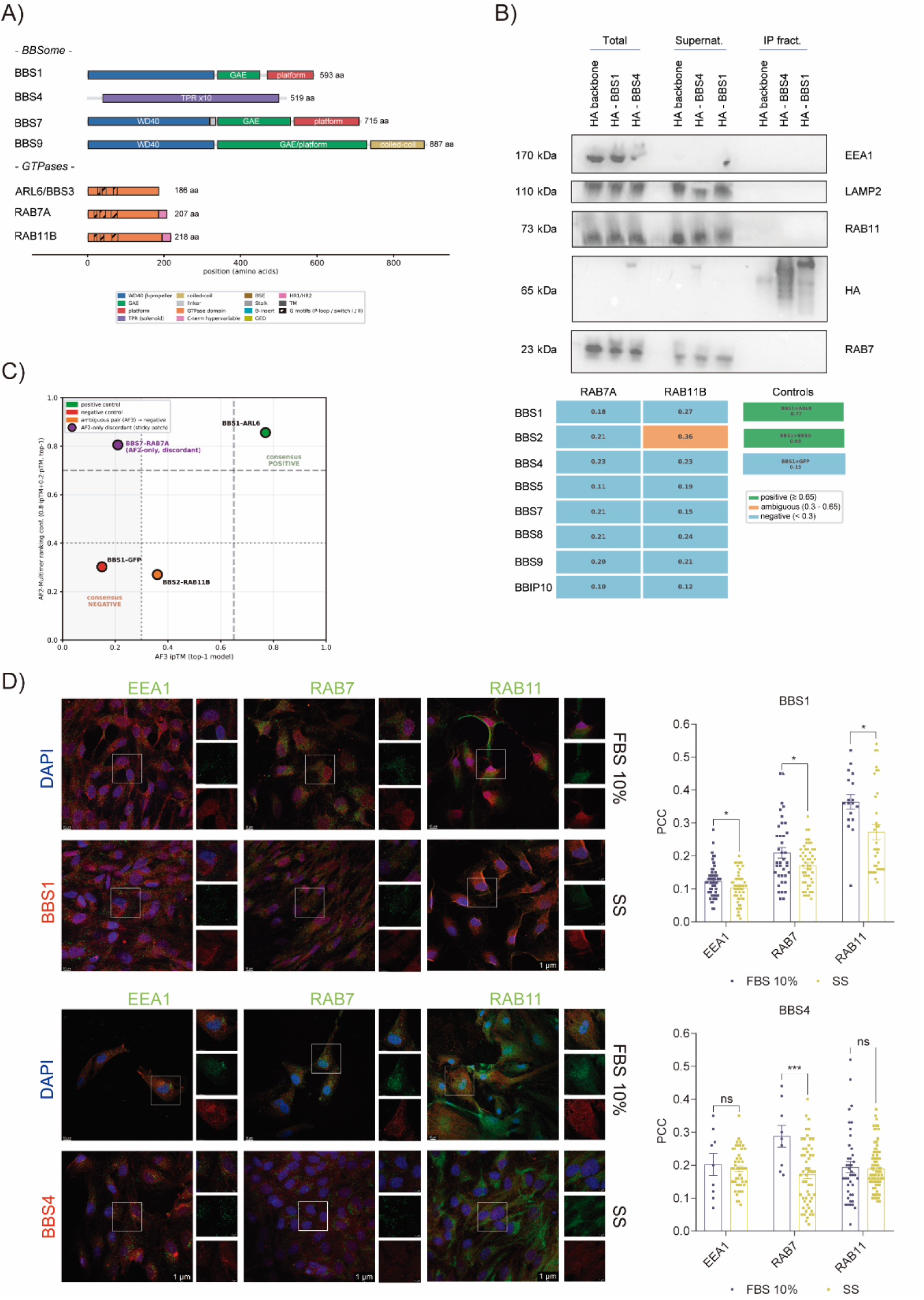
**BBS1 and BBS4 do not form stable complexes with the endocytic trafficking machinery**. (A) Schematic diagram of the BBS protein domain architecture and the GTPases RAB7A, RAB11B, and ARL6, highlighting the structural domains used to guide AlphaFold-based structural modelling. (B) Co-immunoprecipitation of HA-BBS1 and HA-BBS4 overexpressed in HEK293T cells, with the total protein fraction, supernatant and immunoprecipitated fraction probed for EEA1, RAB11, RAB7, LAMP2, and HA, alongside a matrix of AlphaFold 3 ipTM values for pairwise interactions between BBSome subunits and the GTPases RAB7A and RAB11B, calibrated against BBS1–ARL6 (positive; ipTM 0.77), BBS1–BBS9 (positive; ipTM 0.69) and BBS1–GFP (negative; ipTM 0.15) controls. AlphaFold 3 predictions: 5 diffusion samples × 10 recycles per pair; ipTM values shown for the top-ranked model. (C) Cross-engine comparison of selected control and ambiguous pairs, showing AlphaFold 3 ipTM against AlphaFold 2-Multimer ranking confidence scores (0.8·ipTM + 0.2·pTM for the top-ranked model). The panel presents selected validation examples and one AF2-high/AF3-low discordant case; it does not represent a dataset-wide calibration. The borderline AlphaFold 3 pair BBS2–RAB11B was not supported by AlphaFold 2-Multimer (ranking confidence 0.27), whereas BBS7–RAB7A showed high AlphaFold 2-Multimer confidence in 2 of 5 models but lacked support from AlphaFold 3 and manual interface inspection, consistent with a non-specific sticky-patch artefact on the BBS7 GAE+platform region. Co-immunoprecipitation panels are representative of n = 1 independent experiments using clones 1 and 2 of each KO genotype. (D) Immunofluorescence analysis and quantification of colocalisation, measured by the Pearson correlation coefficient (PCC), of BBS1 and BBS4 proteins with EEA1 for early endosomes, RAB11 for recycling endosomes, and RAB7 for late endosomes, providing information on the roles of BBS1 and BBS4 in different stages of the endocytic pathway. The error bars indicate the standard error of the mean (SEM). The number of samples analysed is n = 50. The differences were considered significant when ∗p < 0.05, ∗∗p < 0.01, ∗∗∗p < 0.001.

HA-BBS1 and HA-BBS4 were both efficiently immunoprecipitated, as expected, and all four vesicle markers were detected at the correct size in the total protein fraction and in the supernatant, confirming the assay was technically sound. However, none of the four markers was detected in the immunoprecipitated fraction beyond HA itself (Figure 2B), indicating that BBS1 and BBS4 do not engage in stable, detergent-resistant complexes with EEA1, RAB11, RAB7 or LAMP2. To complement these biochemical data, we used AlphaFold 3 to predict pairwise interactions between BBSome subunits and the GTPases RAB7A and RAB11B, calibrated against three controls: BBS1–ARL6 (a confirmed positive interaction, PDB 4V0O; ipTM 0.77), BBS1–BBS9 (a positive intra-BBSome contact; ipTM 0.69), and BBS1–GFP (negative; ipTM 0.15). Neither BBS1 nor BBS4 showed a positive interaction with RAB7A or RAB11B (ipTM 0.18–0.27 across all four pairs, well below the 0.65 threshold and close to the negative control); BBS1 retained a high-confidence interaction with ARL6 specifically, indicating that its GTPase-binding site is not generic to the RAB7A/RAB11B candidates tested, and BBS4 showed no binding site for either GTPase. Extending the screen to the remaining BBSome subunits identified no positive pairs either: one borderline case (BBS2–RAB11B, AlphaFold 3 ipTM 0.36) resolved to negative on cross-validation with AlphaFold 2-Multimer (ranking confidence 0.27), and an apparent AlphaFold 3/AlphaFold 2 discrepancy for BBS7– RAB7A was traced to a non-specific, recurrent contact surface on BBS7 that does not correspond to its documented binding interface (Figure 2B, C). For BBS1 and BBS4 specifically, then, neither co-IP nor structural modelling supports stable, direct binding to EEA1, RAB11, RAB7 or LAMP2; we do not interpret this as ruling out engagement with other endocytic components, or interfaces involving multiple BBSome subunits simultaneously, which neither assay was designed to detect. Altered TGFBR1 endocytic dynamics and hyperactivation of canonical TGF-β signalling in *BBS1* and *BBS4* KO cells. To further explore whether BBS1 and BBS4 associate with endocytic vesicles in cells, we performed IFM analysis in RPE1 WT cells and quantified colocalization with markers of the endocytic pathway using the Pearson correlation coefficient (PCC). We assessed EEA1, RAB11 and RAB7 under both full-growth and serum-depleted conditions (Figure 2D). This analysis showed that BBS1 colocalized most strongly with RAB11 in both culture conditions, while its overall colocalization with endocytic markers decreased under serum deprivation. In contrast, BBS4 showed a significant increase in colocalization only with RAB7 under serum-supplemented conditions, whereas its association with the other markers changed minimally under serum depletion. Together, these data suggest that BBS1 and BBS4 may associate with distinct endocytic vesicles in a context-dependent manner, but do not support stable direct binding to the limited set of markers tested.

Although our biochemical and structural data argue against stable, direct binding between BBS1/BBS4 and the core endocytic markers tested, the BBSome could still influence the kinetics of receptor trafficking indirectly. To investigate possible alterations in the endocytic dynamics of TGFBR1 in the KO cell lines, we performed colocalization experiments with the same four endosomal/lysosomal markers used throughout this study (Figure 3). Cells were serum-starved for 24 h to enhance ciliogenesis and then stimulated with TGF-β, with images acquired at 0, 5, 10, 15, 30, and 60 min post-stimulation (Figure 3A). Mechanisms of endocytosis in quiescent cells are not fully understood but are believed to be essential for key cellular functions, such as primary cilia formation, cell polarisation, and regulation of cell-cell junctions (41,42), all of which are related to our findings on the proteome.

**Figure 3.**
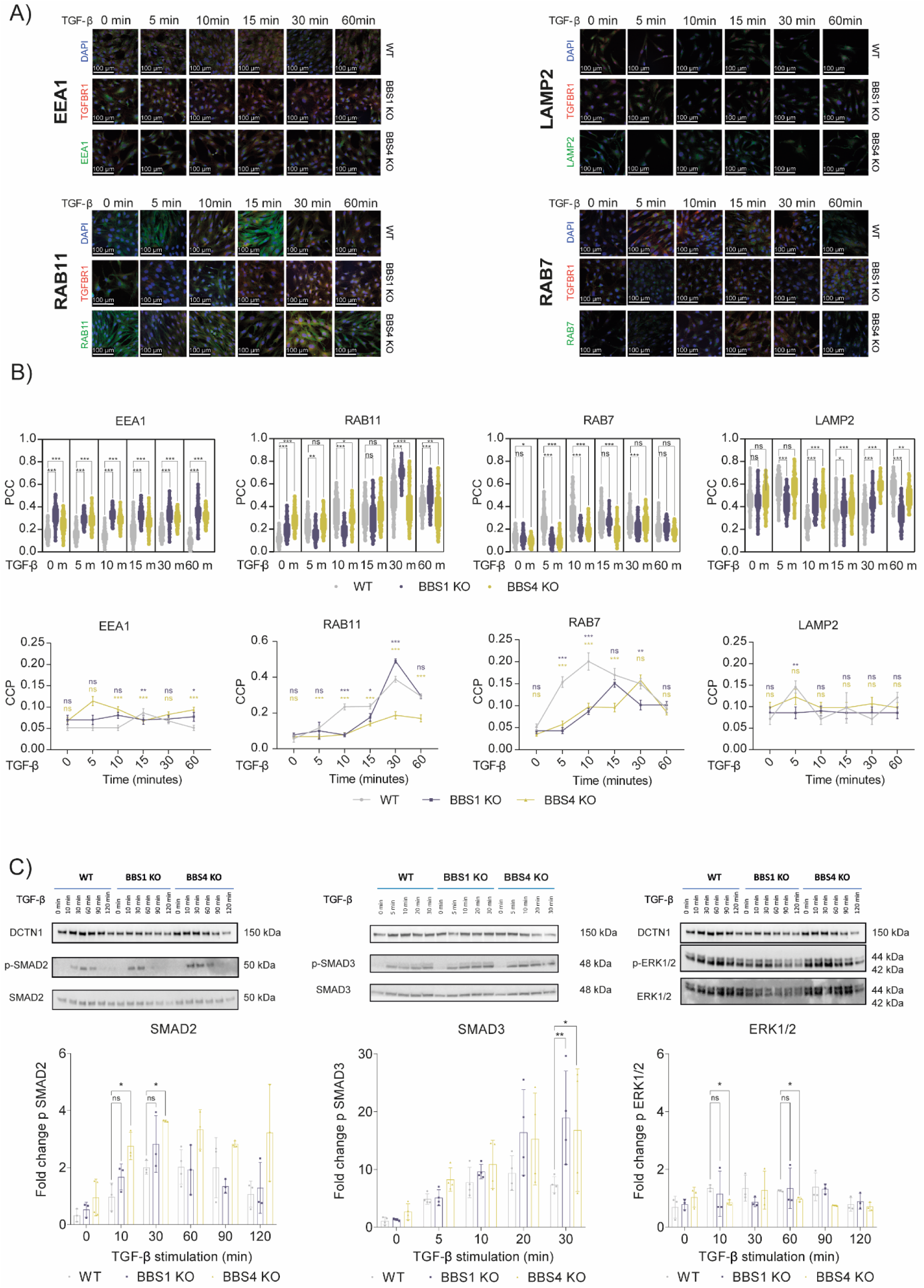
Endocytic dynamics of TGFBR1 and downstream pathway activation in KO cell lines. (A) Immunofluorescence colocalisation of TGFBR1 (594 nm) with EEA1, RAB11, RAB7, and LAMP2 (488 nm), with DAPI counterstain, in WT, BBS1 KO and BBS4 KO cells following 24 h of serum starvation and TGF-β stimulation (0, 5, 10, 15, 30, and 60 min). Scale bar: 100 µm. (B) Quantification of colocalisation by Pearson’s correlation coefficient (PCC), shown as raw values relative to WT and as values normalised to the average WT signal at each time point. (C) Western blot analysis of phosphorylated SMAD2, SMAD3 (canonical pathway) and ERK1/2 (non-canonical pathway) at the same time points. Data represent mean ± SEM from n = 3 independent biological replicates using all clones available for each KO genotype. For colocalisation quantification (B), 50 cells per condition per replicate were analysed. The differences were considered significant when *p < 0.05, **p < 0.01, ***p < 0.001, and ****p < 0.0001. ns = non-significant.

Considered as raw differences in Pearson’s correlation coefficient (PCC) relative to WT, both KO lines showed higher colocalization of TGFBR1 with EEA1 at every time point analysed. Colocalization with RAB11 was likewise higher than WT in both KO lines at all time points except 15 min, with *BBS4* KO additionally not differing from WT at 5 min. The opposite pattern was observed for RAB7, where both KO lines showed significantly lower colocalization than WT at several early-to-intermediate time points (5, 10, and 30 min for *BBS1* KO; 0, 5, 10, and 15 min for *BBS4* KO). For LAMP2, both KO lines were higher than WT at 10, 15, and 30 min but lower at 60 min, with *BBS1* KO also lower at 5 min (Figure 3B).

To better capture the temporal dynamics of TGFBR1 trafficking, we next normalised PCC values against the average WT signal at each time point. For EEA1, the WT response peaked at 15 min before gradually declining, whereas both KO lines already showed (non-significantly) higher colocalization at time 0, peaking earlier (at 5 min in *BBS4* KO and 10 min in *BBS1* KO) and remaining more sustained over time; only at 15 min did WT colocalization significantly exceed that of the KO lines, and by 60 min both KOs were again significantly above WT. RAB11 dynamics followed a broadly similar shape, with a gradual increase to a peak at 30 min followed by decline, but with marker-specific differences between the two KO lines: *BBS4* KO colocalization with RAB11 remained significantly lower than WT across the entire time-course, while *BBS1* KO showed a more abrupt response, with significantly lower colocalization at 10 and 15 min but significantly higher colocalization than WT at the 30-min peak. RAB7 dynamics differed substantially between WT and the KO lines: both KOs showed a gradual rise to a peak (15 min in *BBS1* KO, 30 min in *BBS4* KO) followed by decline, whereas WT colocalization was elevated throughout the entire time-course, rising rapidly from as early as 5 min, peaking at 10 min and then gradually declining; notably, the WT value at 15 min coincided with the *BBS1* KO peak, and the WT value at 30 min coincided with the *BBS4* KO peak. LAMP2 dynamics were the most divergent of the four markers: WT cells displayed cyclical colocalization peaks approximately every 5 min, the largest occurring 5 min after stimulation; *BBS4* KO showed weaker cyclical peaks roughly every 10 min; and *BBS1* KO showed essentially no temporal response, with colocalization values remaining well below those of the other two cell lines throughout (Figure 3B).

This pattern of kinetic differences indicates that both *BBS1* and *BBS4* deficiency delay the resolution of TGFBR1 from early endosomes and alter its partitioning between the recycling (RAB11) and degradative (RAB7/LAMP2) arms of the endocytic pathway, but with distinct dynamics for each subunit. *BBS1* KO cells show an exaggerated, delayed recycling response, reaching their highest relative RAB11 colocalization at the time WT cells are already favouring degradation, together with an essentially absent late lysosomal response. *BBS4* KO cells, in contrast, show persistently reduced RAB11 engagement across the full time-course, accompanied by a blunted, but not absent, lysosomal response.

To determine whether these trafficking alterations translate into changes in TGF-β pathway activation, we examined phosphorylation of SMAD2/3 (canonical pathway) and ERK1/2 (non-canonical pathway) by Western blot at the same time points (Figure 3C). Both KO lines showed overexpression of the canonical pathway at 30 min, as indicated by increased pSMAD3, with this effect markedly stronger in *BBS4* KO cells, which also showed increased pSMAD2 at 10 and 30 min. In contrast, activation of the non-canonical ERK1/2 pathway was reduced in *BBS4* KO cells at 10 and 60 min. These results indicate that, despite their distinct TGFBR1 trafficking signatures, both KO lines converge on hyperactivation of canonical TGF- β/SMAD signalling, with *BBS4* deficiency associated with the more pronounced effect.

### BBS1 and BBS4 deficiency led to distinct, opposing alterations in autophagic flux

TGFBR1 can be degraded via ubiquitination or through the lysosomal pathway, depending on the proteins involved (43,44). For example, SMAD7, which in quiescent cells localises to the CiPo area (45,46), binds to SMURF2 to form an E3 ubiquitin ligase that targets TGFBR1 for degradation (47). Given the differential lysosomal (LAMP2) engagement of TGFBR1 we observed between *BBS1* KO and *BBS4* KO cells (Figure 3), and the central role of autophagy in RPE1 cells for recycling defective photoreceptor components, we next examined whether these trafficking differences were accompanied by alterations in autophagic flux.

We performed Western blot analysis of p62, TGFBR1 and LC3B (forms I and II) in WT, *BBS1* KO and *BBS4* KO cells following TGF-β stimulation (0, 30, and 60 min), with a 60 min Bafilomycin A1 (BafA1) condition included to block autophagosome–lysosome fusion (Figure 4). TGFBR1 was present at significantly higher levels in both KO lines than in WT, consistent with our proteomic data for *BBS1* KO (Figure 1) and extending this observation to *BBS4* KO. p62 levels changed significantly over time only in *BBS1* KO cells, where they decreased progressively with stimulation; TGFBR1 followed a similar declining trend in this line. Notably, the decrease in TGFBR1 in *BBS1* KO cells remained significant even in the presence of BafA1, indicating that this decline is not fully autophagy-dependent and may involve additional degradative routes, such as direct lysosomal degradation following endocytosis or proteasomal degradation.

**Figure 4.**
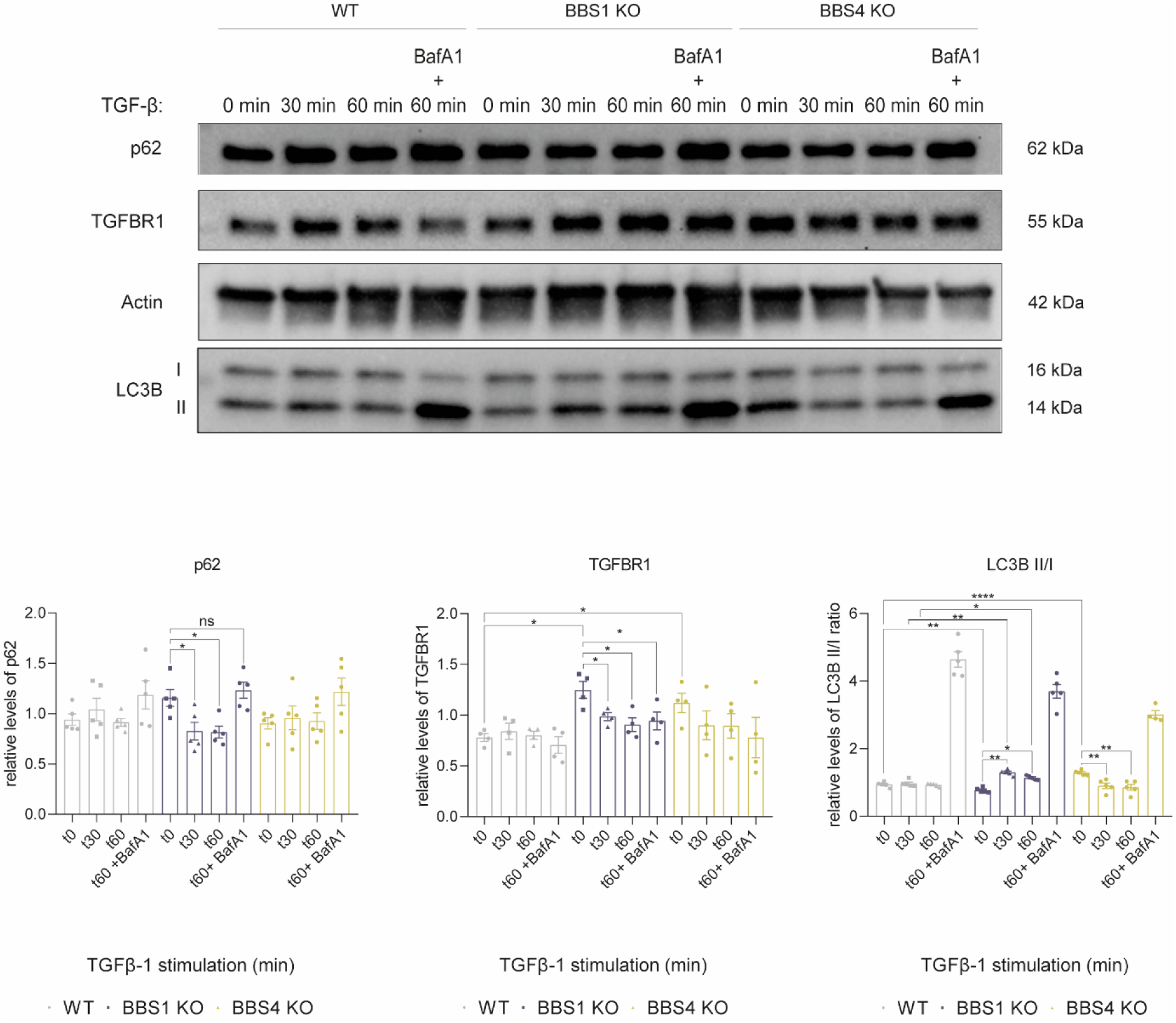
**Differential autophagic flux in BBS1 and BBS4 KO cells following TGF-β stimulation**. Western blot analysis of p62, TGFBR1 and LC3B (forms I and II) in WT, BBS1 KO, and BBS4 KO cells at 0, 30, and 60 min of TGF-β stimulation following 48 h of serum starvation, with an additional 60 min Bafilomycin A1 (BafA1, 20 nM; autophagy inhibitor) condition included as a degradative flux control. The LC3B-II/I ratio was calculated as a measure of autophagosome abundance; all signals were normalised to actin. Data represent mean ± SEM from n = 5 independent biological replicates using all clones available for each KO genotype. Significant differences in the LC3B-II/I ratio relative to t = 0 and the BafA1 condition are not annotated on the figure due to space constraints but are described in the Results. The differences were considered significant when *p < 0.05, **p < 0.01, ***p < 0.001, and ****p < 0.0001. ns = non-significant.

The most pronounced differences were observed in the LC3B-II/I ratio, a measure of autophagosome abundance. Under basal, unstimulated conditions, this ratio was significantly higher in *BBS4* KO and significantly lower in *BBS1* KO than in WT. Upon TGF-β stimulation, the ratio increased significantly in *BBS1* KO at 30 min and remained elevated, albeit to a lesser extent, at 60 min. In the presence of BafA1, both KO lines showed significantly lower LC3B- II/I ratios than WT.

Within each cell line, WT cells showed a significant change only in the presence of BafA1, where the LC3B-II/I ratio increased markedly, as expected for cells with normal, unperturbed autophagic flux. *BBS1* KO cells, by contrast, showed a significant increase in the ratio upon TGF-β stimulation at 30 min (and, to a lesser extent, at 60 min) relative to unstimulated cells, in addition to the expected BafA1-induced increase. *BBS4* KO cells showed the opposite pattern: a progressive and significant decrease in the LC3B-II/I ratio during stimulation, followed by a sharp increase upon BafA1 treatment, indicating that, despite this decline, the autophagic machinery in *BBS4* KO cells remains inducible and the BafA1 response itself is intact.

These results revise our earlier assessment, in which we found no significant differences in autophagy markers between KO and WT cells; the present time-resolved analysis instead indicates that *BBS1* and *BBS4* deficiency are each associated with distinct, and largely opposite, alterations in autophagic flux upon TGF-β stimulation. *BBS1* KO cells, which show low basal autophagosome abundance, mount an induced autophagic response upon stimulation; *BBS4* KO cells, which show elevated basal autophagosome abundance, show a stimulation-dependent decline in steady-state LC3B-II/I that is nonetheless reversible by blocking degradative flux, consistent with retained but more rapidly turned-over autophagosomes. Proteins directly linked to autophagy may also act outside strict degradative roles: Beclin-1, for example, is required for recycling of TGFBR1, facilitating its localisation to RAB11+ endosomes (48), raising the possibility that altered autophagy-related protein activity contributes to the distinct TGFBR1 trafficking itineraries described above (Figure 3), rather than acting solely as a degradative endpoint.

### Loss of BBS proteins enhances cell migration and promotes an EMT phenotype

Given that both *BBS1* and *BBS4* KO cells showed hyperactivation of canonical TGF-β/SMAD signalling downstream of altered TGFBR1 trafficking (Figure 3C), together with distinct, opposing alterations in autophagic flux (Figure 4), we next examined the functional consequences for EMT, a TGF-β-driven process of particular relevance to RPE biology: quiescent RPE cells are integrated into tissue via cell-cell junctions at their basolateral membrane, and these junctions are characteristically lost during EMT.

We first examined EMT markers by immunofluorescence in unstimulated cells and after 6 h of TGF-β stimulation, using N-cadherin and ACTA2 as mesenchymal markers (Figure 5A). Although differences did not reach statistical significance, both KO lines showed a trend towards accumulation of both markers relative to WT, both at baseline and following stimulation. As previously observed, in *BBS1* KO cells, ACTA2 filament formation was already detectable at 0 h, whereas in *BBS4* KO cells, filaments were absent at 0 h, with ACTA2 instead concentrated in a perinuclear, crescent-shaped pattern, only forming filaments after stimulation.

**Figure 5.**
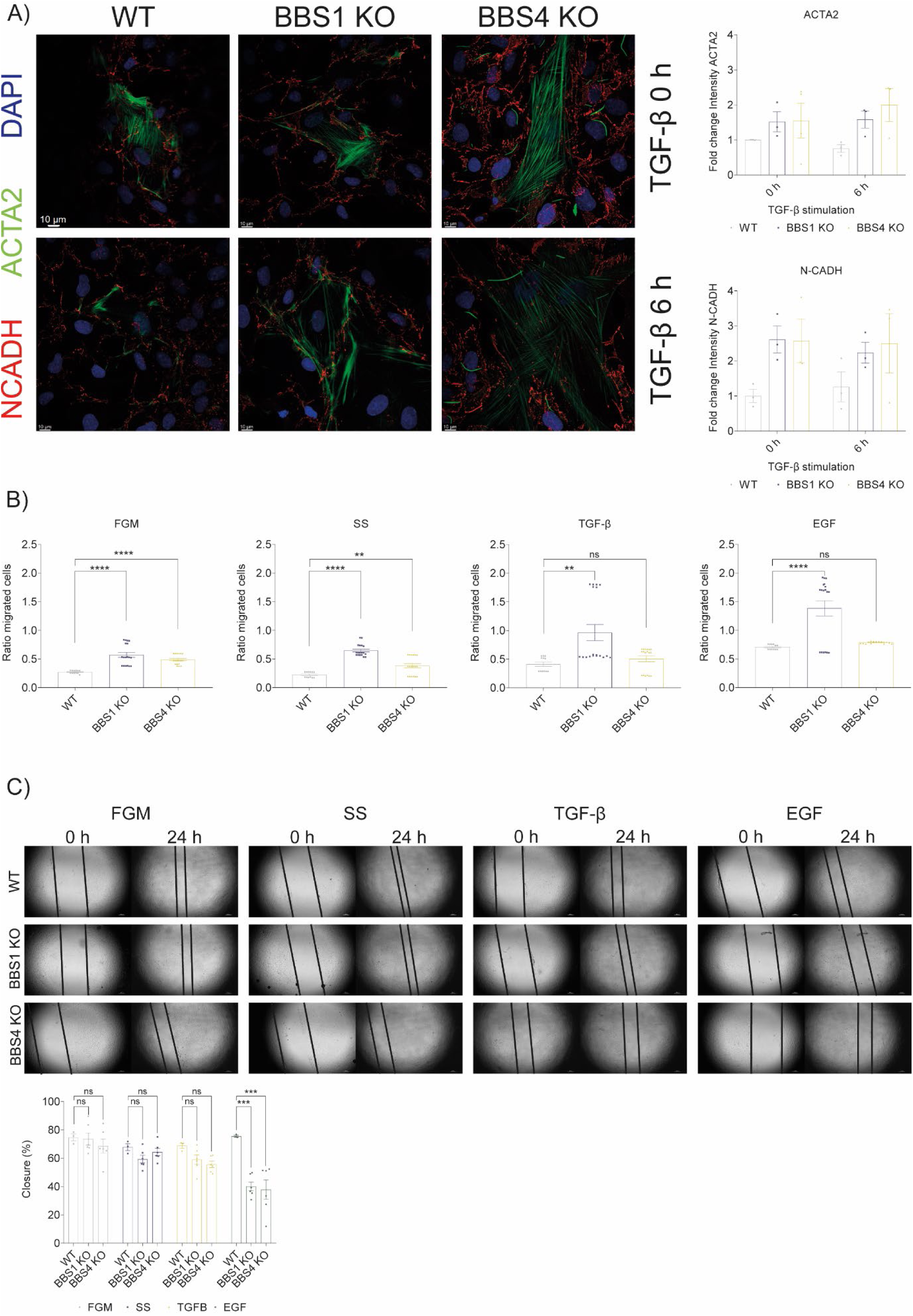
EMT assays in KO cell lines. (A) Immunofluorescence of N-cadherin (594 nm) and ACTA2 (488 nm) as EMT markers, with DAPI counterstain, in WT, BBS1 KO, and BBS4 KO cells in unstimulated conditions and after 6 h of TGF-β stimulation. Scale bar: 10 µm. Staining intensity was quantified from 4 fields of view per condition. (B) Transwell migration assay under full growth medium (FGM), serum-starved medium (SS), TGF-β-supplemented medium (2 ng/mL), and EGF-supplemented medium (20 ng/mL) as a positive control. (C) Wound- healing assay under the same panel of conditions. For (B) and (C), data represent mean ± SEM; n = 6 inserts per condition across 3 independent biological replicates. The differences were considered significant when *p < 0.05, **p < 0.01, ***p < 0.001, and ****p < 0.0001. ns = non-significant.

To test whether this mesenchymal predisposition translated into altered migratory behaviour, we performed a transwell assay across complete growth medium, serum-deprived medium, and TGF-β-supplemented medium, with EGF as a positive control (Figure 5B). Both KO lines showed significantly higher migratory capacity than WT across all conditions tested: *BBS1* KO cells nearly doubled their migratory capacity compared to WT, while *BBS4* KO cells also showed increased, albeit comparatively smaller, migration. Under EGF stimulation, the *BBS1* KO line migrated furthest of all three cell lines.

We complemented this with an *in vitro* wound-healing assay under the same panel of treatments (Figure 5C). The EGF positive control confirmed that both KO lines have a reduced capacity for wound closure relative to WT; however, no significant differences were observed across the other treatment conditions, with only a trend towards reduced wound-healing capacity detected in the KO lines.

To confirm these phenotypic observations at the molecular level, we measured the expression of a panel of epithelial and mesenchymal markers by qPCR, including several identified by our proteomic analysis, both under normal growth conditions and following TGF-β stimulation after serum starvation (Figure 6A). Under normal culture conditions, *BBS1* KO cells already showed a mesenchymal predisposition, with significant activation of EMT markers such as *CDH2* and downregulation of markers such as *CDH1*. The results on *SNAI1* and *SNAI2* or *TWIST1* transcription factors that activate EMT processes contrast, and in *BBS1* KO they are not active or even appear repressed, even under stimulation with TGF-β. In contrast, *BBS4* KO cells presented a more epithelial phenotype, with reduced expression of *CDH2*, *ACTA2*, and *CRYAB*. However, after 48 h of serum depletion and 30 min of TGF-β stimulation, the EMT activity increased. *BBS1* KO cells exhibited a slight but significant increase in *ACTA2* and *CRYAB* expression, along with a notable decrease in *CDH1* expression. *BBS4* KO cells also showed decreased *CDH1* levels, but increased expression of *CDH2*, *VIM*, and *SNAI1*. *CTNNB1* expression increased significantly in both cell lines after stimulation.

**Figure 6.**
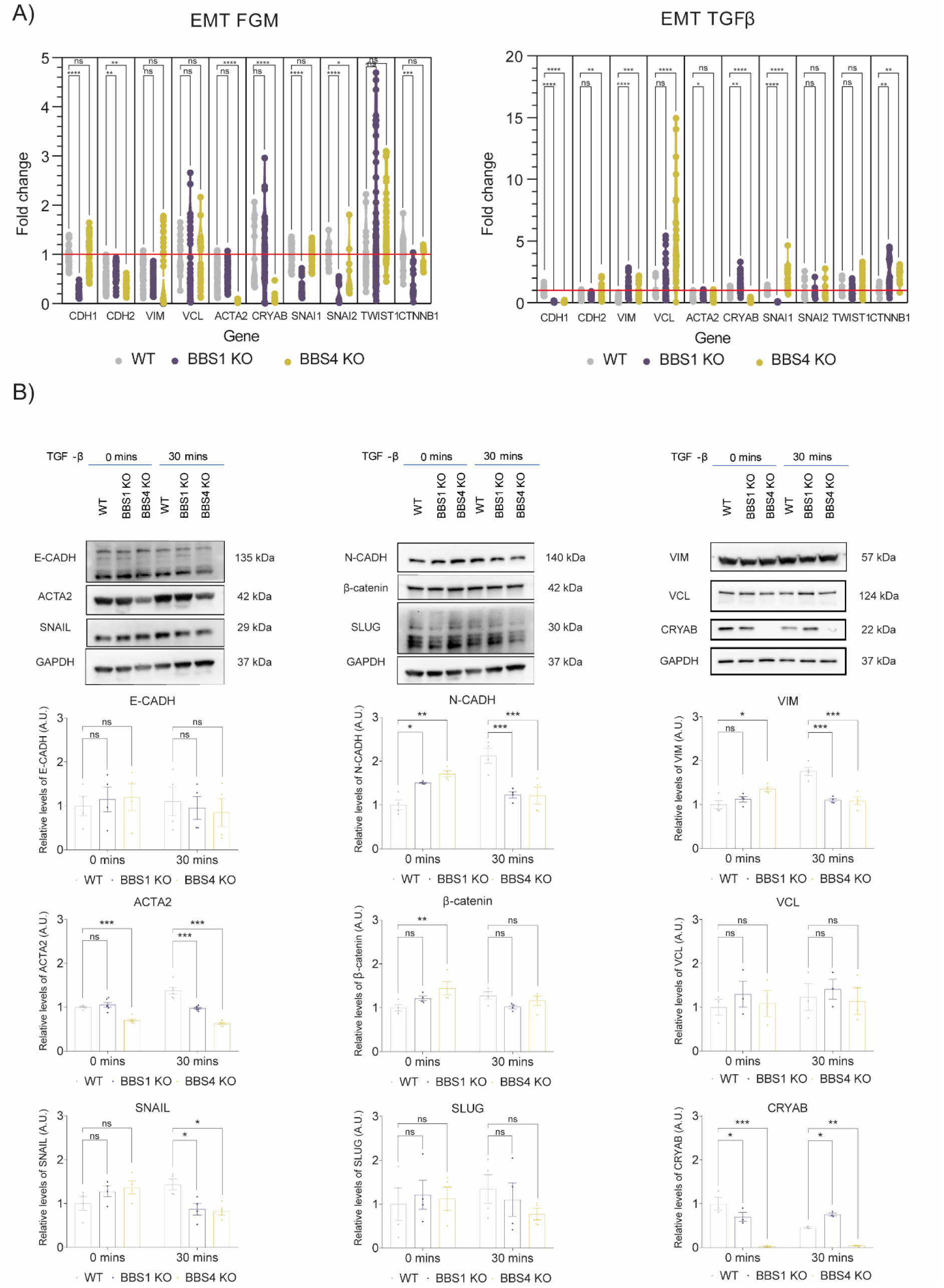
EMT marker expression in KO cell lines. (A) qPCR analysis of a panel of epithelial and mesenchymal markers, including several identified by surface proteomic analysis (Figure 1), under normal growth conditions and following TGF-β stimulation after 48 h of serum starvation. Each data point represents an independent biological replicate; each replicate was measured in technical triplicate and normalised to WT. (B) Western blot analysis of the same marker panel at 0 and 30 min of TGF-β stimulation, with actin as loading control. Representative blots are shown. Data represent mean ± SEM from n = 4-15 independent biological replicates (qPCR) and n = 4 independent biological replicates of all clones available for each KO genotype throughout. The differences were considered significant when *p < 0.05, **p < 0.01 and ***p < 0.001. ns = non-significant.

To confirm the mesenchymal phenotype observed in the KO lines, particularly in *BBS1* KO, we performed Western blot analyses of the same panel of EMT markers assessed by qPCR, measured at baseline and 30 min after TGF-β stimulation, the time at which canonical SMAD2/3 activation peaks in our quiescent RPE1 cells (Figure 6B). Both KO lines exhibited a predisposition toward an EMT phenotype, confirmed by qPCR and WB for the N-cadherin and β-catenin markers, where higher expression was observed at time 0 min in KO cells, followed by a decrease at 30 min compared to WT. SNAIL and SLUG levels, although not significant, are higher in both KO cell lines at time 0, remain stable or decrease slightly at 30 min, while they increase in WT cells. This is in line with what was postulated for N-cadherin levels: KO cells start from a more mesenchymal stage, and the pathway is saturated earlier than in WT cells. Although ACTA2 baseline levels were consistent with qPCR and IFM data, at 30 min, WB showed a decrease that did not align with the qPCR results. IFM at 6 h revealed the highest filament formation in KO cells, suggesting that ACTA2 protein levels may respond more slowly than transcript levels. VIM also showed a significant reduction at 30 min, indicating variability in marker response times. However, the most noteworthy case is that of CRYAB, which aligns perfectly with the gene expression results: it is undetectable in *BBS4* KO but significantly accumulates in *BBS1* KO following stimulation. Studies have shown that αβ-crystallin modulates EMT through SNAIL and SLUG, acting as a molecular chaperone for SMAD4 (49). These findings in human RPE1 cells are consistent with the EMT-like traits previously observed in Bbs8-deficient mice and RPE in vivo (50), and extend them to BBS1 and BBS4 deficiency in a human cell model.

## Discussion

Our study aimed to explore the role of *BBS1* and *BBS4* KO retinal cells using plasma membrane proteomics to better understand the molecular mechanisms of BBS, particularly in receptor endocytosis. Retinal degeneration affects nearly 100 % of people with BBS, characterised by early loss of photoreceptors, night blindness, and progressive visual decline leading to blindness (6,51). Mouse models show that despite photoreceptor degeneration, some functionality remains, which highlights the importance of understanding the tissue-specific regulation of BBSome entry into cilia (52).

Given the importance of RPE integrity for photoreceptor health, we used hTERT-RPE1 cells to model the KO phenotypes of *BBS1* and *BBS4*. Both KO lines showed shorter cilia, but with different ciliation frequencies: *BBS1* KO cells retained normal ciliation rates with shorter cilia, while *BBS4* KO cells showed fewer cilia. This suggests distinct roles for BBS1 and BBS4, consistent with their roles in pre-BBSome nucleation and BBSome assembly (27). BBS4 helps nucleate the pre-BBSome in the pericentriolar satellites, while BBS1 mediates its transport to the ciliary base, showing that both are essential for proper BBSome function in the primary cilia.

Ciliary dysfunction is associated with impaired endocytosis. The BBSome seems essential for correctly internalising receptors at the CiPo and recycling back to the plasma membrane, at least in invertebrates, where cilia-independent functions are already described: *T. brucei* BBS proteins form a BBSome that interacts with clathrin and is located on the membranes of the flagellar pocket and nearby cytoplasmic vesicles (53), and in the *C. elegans* photoreceptor protein LITE-1 stability in ciliated ASH photosensory neurones is regulated by BBSome through Rab5-mediated endocytosis (54). Notably, our co-immunoprecipitation and AlphaFold analyses did not support an equivalent stable, direct association between BBS1/BBS4 and the core endocytic markers examined in RPE1 cells, suggesting that this mode of BBSome engagement with the endocytic machinery may not directly transfer to mammalian, cilia- bearing epithelial cells, or that it relies on partners other than EEA1, RAB11, RAB7 or LAMP2. Several signalling pathways are involved in RPE differentiation, including ciliary pathways such as Shh, Wnt/β-catenin, Notch, and TGF-β (55–57). Studies in zebrafish and human cells have shown that BBSome plays a crucial role in internalising receptors, such as leptin and insulin receptors, and recycling them to the plasma membrane (16,18,19). Mislocalization and impaired trafficking of these receptors, particularly leptin and insulin receptors, contribute to the metabolic dysfunctions seen in BBS, such as hyperphagia and obesity (58).

Endocytosis regulates receptor availability on the cell surface, with continuous internalisation preventing excessive signalling and cell cycle re-entry (59). This regulation is essential for the TGF-β pathway, where normal cells degrade internalised TGFBR1 to limit signalling and prevent EMT. The BBSome has structural similarities to coat complexes such as COPI, COPII, and clathrin, hinting at a shared evolutionary origin (60–62). TGF-β receptor internalisation involves clathrin- and caveolin-mediated endocytosis; CME supports receptor recycling and signalling, while caveolin-mediated endocytosis facilitates signal termination and degradation (63,64). Research suggests a convergence of these pathways, where vesicles can form with both caveolin-1 (CAV1) and clathrin, potentially enabling receptors internalised through caveolin-mediated endocytosis to activate R-SMADs similarly to the clathrin-mediated pathway (65).

In RPE1 cells, *BBS1* and *BBS4* KO lines show distinct, but overlapping, alterations in TGFBR1 trafficking. *BBS1* KO cells show sustained early-endosome residency together with an exaggerated, delayed peak in recycling-endosome (RAB11) engagement, consistent with receptor recycling outpacing degradation and driving a robust mesenchymal transcriptional and protein signature. *BBS4* KO cells, in contrast, show persistently reduced recycling-endosome engagement across the full time-course, yet still show hyperactivation of canonical SMAD signalling, even more pronounced than in *BBS1* KO, together with reduced activation of the non-canonical ERK1/2 branch. This indicates that defective BBSome function need not act through a simple recycling-versus-degradation switch to dysregulate TGF-β signalling, and that the relative balance between canonical and non-canonical pathway activation, rather than receptor fate alone, may determine the extent to which hyperactivated signalling is translated into a complete EMT phenotype, a possible explanation for the comparatively milder transcriptional and protein-level EMT signature observed in *BBS4* KO cells despite their stronger SMAD activation.

Assembly of the BBSome begins at the pericentriolar satellites, nucleated by BBS4, and proceeds via BBS1-mediated transport to the ciliary base, suggesting that loss of BBS4 could compromise BBSome assembly more broadly than loss of BBS1 (27). This more global impairment may explain why *BBS4* KO cells show a more uniformly reduced engagement with the recycling pathway across the full time-course, together with a blunted, but not absent, lysosomal response, in contrast to *BBS1* KO cells, which show an essentially absent late degradative response but an exaggerated, delayed recycling peak. Overactivation of the R-SMAD pathway in both KOs could result from pathway crossover in double-positive EEA1 and CAV1 endosomes (65).

The opposing autophagic phenotypes we observed add a further layer to this picture. *BBS1* KO cells, which showed an essentially absent late lysosomal (LAMP2) response in our TGFBR1 trafficking time-course (Figure 3), nonetheless mount a clear stimulation-induced increase in autophagosome abundance (Figure 4), suggesting that reduced LAMP2 engagement in this line does not reflect a general degradative deficiency, but may instead be specific to the canonical endolysosomal route, with autophagy partially compensating. *BBS4* KO cells, conversely, combine elevated basal autophagosome abundance with a stimulation-induced decline that is reversible by blocking flux, consistent with retained but accelerated autophagic turnover rather than a block in the pathway. Whether this more dynamic autophagic response in *BBS4* KO contributes to its stronger canonical SMAD activation (Figure 3C), for instance, via autophagy-dependent turnover of signalling intermediates, is an open question that the present data cannot resolve, but one we consider worth pursuing given the established role of selective autophagy in shaping receptor tyrosine kinase and TGF-β signalling duration.

Our proteomic analysis found a significant enrichment of TGFBR1 on the plasma membrane of BBS KO cells, which influences EMT through R-SMAD signalling, which depends on CME. Our findings on altered endocytosis in KO cells suggest that BBSome function differs between receptor types, playing a key role in the coordination of cilia-dependent and cilia-independent endocytic pathways. TGF-β signalling, closely linked to receptor internalisation, requires CME for proper activation (66–68). Upon ligand binding, TGFBR1 is internalised via the ciliary pocket to become integrated with early endosomes, which is the site for robust R-SMAD activation (12). The fate of the receptor is then determined: RAB7 promotes degradation through late endosomes, while RAB11 supports recycling (69).

In *BBS1* KO cells, we observed an increased accumulation of TGFBR1 in recycling endosomes. This contrasts with a previous study (17), which reported receptor accumulation in late endosomes with reduced recycling. This discrepancy may stem from methodological differences, as we utilised RPE1 KO cells, whereas Leitch et al. employed an siRNA-based knockdown approach. This does not appear to be the only case of a receptor affected by BBS4. Specifically, this laboratory also demonstrates the relationship of this subunit in the expression and secretion of FSTL1, a protein in the TGFB/BMP superfamily (70). The distribution during asymmetric cell division may also play a role in these differences since endosomes containing receptors such as Notch are targeted to one daughter cell during mitosis to regulate cell fate (33,35,71).

In *BBS1* KO cells, markers (e.g., E-cadherin, N-cadherin, VIM, SNAIL, ACTA2) showed a more mesenchymal profile, correlating with increased migration under TGF-β stimulation. This supports the hypothesis that BBSome dysfunction alters receptor trafficking and EMT through R-SMAD signalling, overstimulating the TGF-β pathway. The mislocalization of receptors, as seen with PDGFRα in IFT20-deficient cells, has been associated with excessive signalling (72). Another effect related to disrupted PDGFα receptor signalling is wound healing, as shown in cells and mice with the BBS1^M390R/M390R^ mutation, which exhibits delayed wound closure and impaired migration, consistent with the trend towards reduced wound-healing capacity we observed in our KO lines, most clearly under EGF stimulation (73).

These findings align with previous studies (74), showing the involvement in EMT across different tissues. This study expands the understanding of the role of BBS1 in the regulation of RPE-specific EMT via the TGF-β pathway. Although both *BBS1* and *BBS4* KO cells exhibited altered EMT, BBS1 deficiency led to a more pronounced mesenchymal phenotype and increased migration, potentially disrupting the epithelial barrier vital for photoreceptor support. This contrasts with previous reports suggesting that BBS1 deficiency leads to milder phenotypes (30,31). Given that the p.M390R mutation in *BBS1* is the most common BBS mutation, our results in RPE1 cells could have significant implications for many patients with BBS, particularly concerning gene therapy targeting retinal degeneration. Disruption in the structure and polarity could hinder the effectiveness of gene therapies by compromising the blood-retinal barrier and viral delivery to photoreceptors.

Based on these findings, BBSome regulation of TGFBR1 trafficking does not appear to follow a simple cilia-dependent/independent or recycling/degradation dichotomy. In *BBS1* KO cells, accumulation of TGF-β-pathway proteins on the plasma membrane, together with sustained, delayed recycling-endosome engagement, is associated with robust mesenchymal marker induction and enhanced migration, consistent with excessive receptor recycling driving sustained canonical signalling. In *BBS4* KO cells, persistently reduced recycling-endosome engagement is nonetheless accompanied by the strongest canonical SMAD activation observed in this study, while non-canonical ERK1/2 signalling is reduced; this combination is associated with a comparatively milder and less complete EMT signature than in *BBS1* KO cells, suggesting that the balance between canonical and non-canonical TGF-β signalling, rather than the route of receptor trafficking alone, shapes the extent of EMT execution downstream of BBSome dysfunction. Because our co-immunoprecipitation and AlphaFold analyses do not support stable, direct binding between BBS1/BBS4 and the core endocytic markers examined, we favour a model in which the BBSome influences TGFBR1 trafficking indirectly. For example, through transient or cargo-specific interactions, or through its broader effects on ciliary signalling and cell polarity, rather than by acting as a stable component of the early endosomal, recycling, or degradative machinery itself. We summarise this revised model, integrating receptor partitioning, trafficking, autophagic flux, signalling, and EMT outcomes across genotypes, in Figure 7.

**Figure 7.**
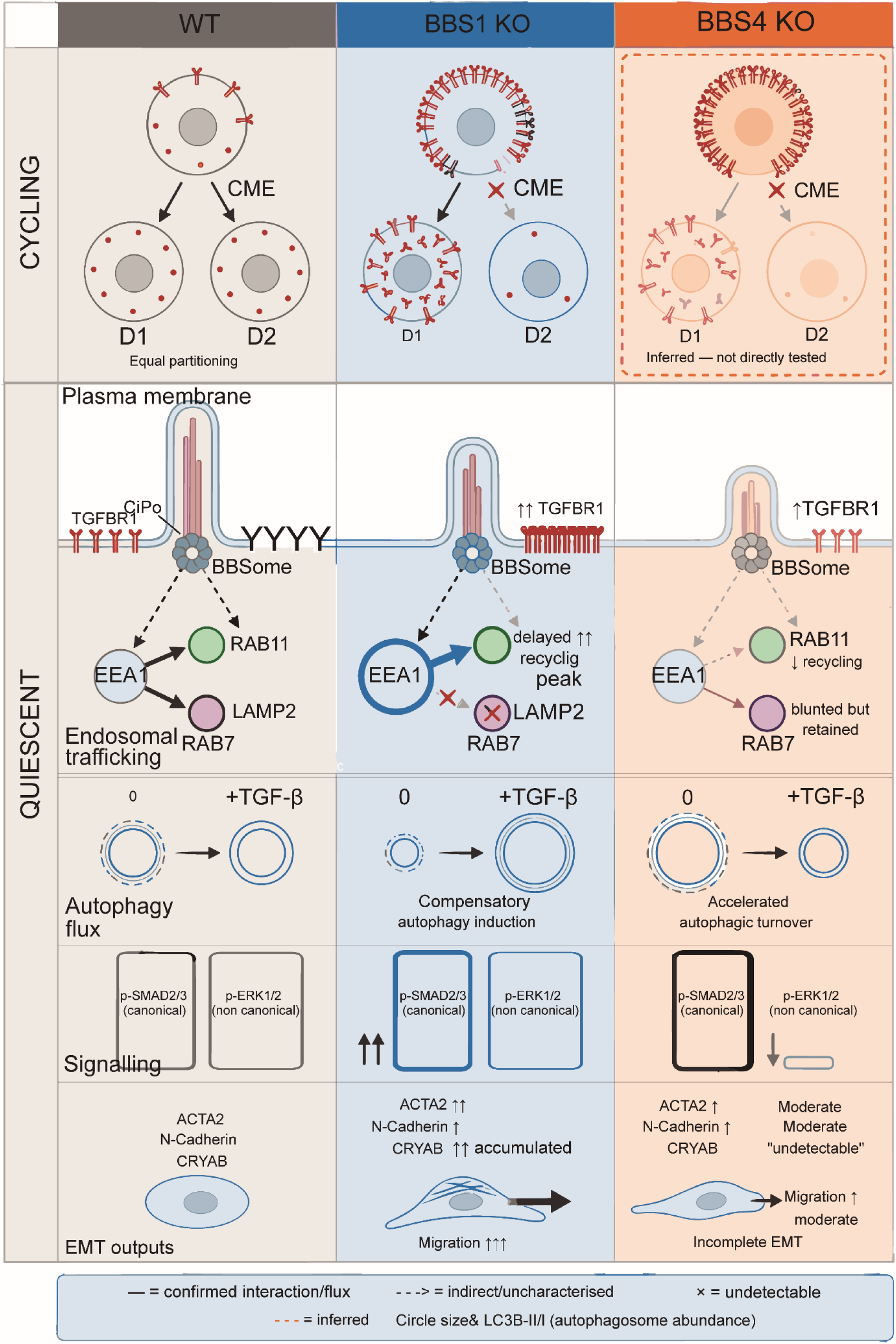
BBSome-dependent TGFBR1 trafficking, autophagic flux, and TGF-β/EMT signalling in RPE1 cells. Genotypes are arranged as columns (WT, BBS1 KO, BBS4 KO) and cell state as rows (cycling; quiescent/ciliated), with the quiescent row subdivided into five mechanistic sub-rows: BBSome and membrane engagement, endosomal trafficking, autophagic flux, downstream signalling, and EMT outputs. In cycling WT cells, CME partitions TGFBR1 equally between daughter cells at mitosis; in BBS1 KO cycling cells, the receptor accumulates at the plasma membrane. The BBS4 KO cycling panel is shown as inferred, as equivalent proteomic evidence was not obtained. In quiescent ciliated cells, the BBSome engages the endocytic machinery indirectly (dashed connectors); no stable physical complex between BBS1/BBS4 and EEA1, RAB11, RAB7 or LAMP2 was detected by co-immunoprecipitation or AlphaFold modelling. BBS1 KO cells show sustained early-endosome retention, an exaggerated and delayed recycling peak, and near-absent lysosomal response, together with low basal autophagosome abundance that increases upon TGF-β stimulation. BBS4 KO cells show reduced recycling-endosome engagement and a blunted lysosomal response, together with elevated basal autophagosome abundance that declines upon stimulation, consistent with accelerated autophagic turnover. Both KO lines show hyperactivation of canonical SMAD2/3 signalling, most pronounced in BBS4 KO cells, which additionally show reduced non-canonical ERK1/2 activation. BBS1 KO cells develop a robust mesenchymal output (CRYAB accumulation, elevated ACTA2/N-cadherin, enhanced migration). BBS4 KO cells, despite stronger canonical signalling, show a mild and incomplete EMT signature with undetectable CRYAB, indicating that the balance between canonical and non-canonical TGF-β signalling, rather than receptor fate alone, determines EMT execution downstream of BBSome dysfunction. CME: clathrin-mediated endocytosis. Dashed connectors: indirect or uncharacterised interaction. ✕: undetectable protein. Circle size proportional to LC3B-II/I ratio. Panel without symbol: inferred.

Our findings provide a mechanistic framework linking BBSome dysfunction to EMT in the RPE. Loss of Bbs8 has been shown to induce EMT-like changes in vivo, including junctional disruption and cytoskeletal reorganisation (50). In parallel, we show that disruption of *BBS1* and *BBS4* also promotes a mesenchymal shift in human RPE cells, but through a defined mechanism involving altered endocytic trafficking of TGFBR1 and differential activation of TGF-β/SMAD and ERK1/2 signalling.

Despite converging on a broadly similar EMT-like phenotype, BBS1 and BBS4 deficiency result in distinct trafficking itineraries, autophagic flux alterations, and signalling outputs. Together with the Bbs8 phenotype, these findings support a model in which BBSome subunit identity shapes the route to TGF-β pathway dysregulation, potentially contributing to variability in EMT-related features across BBS genotypes.

We note that the links proposed here between TGFBR1 trafficking, SMAD/ERK activation and EMT are based on temporal and quantitative correlation across independent assays performed in the same cell lines, rather than on a direct causal or epistasis experiment (for example, pathway-selective inhibition or rescue of trafficking would be needed to establish that the observed trafficking differences are themselves responsible for the differences in EMT output, rather than co-occurring consequences of BBSome loss). We consider this the most important open question raised by the present data, and a priority for follow-up work.

## Conclusions

In conclusion, our study demonstrates that the BBSome, particularly BBS1, is an indirect regulator of receptor endocytosis and signalling, with significant implications for EMT. Notably, this regulation appears to occur independently of stable physical engagement between the BBSome and the core endocytic machinery, pointing towards more dynamic or context-dependent modes of trafficking control that warrant further mechanistic investigation. BBS1 deficiency results in more pronounced EMT alterations in RPE1 cells. The accumulation of TGFBR1 in the plasma membrane and the altered trafficking underscore the broader impact of BBSome dysfunction on receptor dynamics. These findings enhance our understanding of BBS pathology and suggest that targeting the TGF-β pathway may provide new therapeutic strategies for retinal degeneration and other complications associated with BBS (75,76). To this end, further studies should incorporate animal models and patient cells expressing BBS subunit variants to better evaluate the role of the BBSome in EMT processes across different cell types, aiming to develop strategies to slow tissue deterioration. Finally, future research should investigate the potential of combining gene therapy with EMT inhibitors to improve treatment outcomes for BBS patients.

## List of abbreviations

ACTA2: actin alpha 2, smooth muscle
ARL6: ADP-ribosylation factor-like protein 6
BafA1: Bafilomycin A1
BBS: Bardet-Biedl syndrome
BBSome: Bardet-Biedl syndrome octameric protein complex
BME: β-Mercaptoethanol
BMP: bone morphogenetic protein
CiPo: ciliary pocket
CME: clathrin-mediated endocytosis
CRISPR: clustered regularly interspaced short palindromic repeats
CRYAB: crystallin alpha B
DAPI: 4ʹ,6-diamidino-2-phenylindole
DMEM: Dulbecco’s Modified Eagle Medium
EEA1: early endosome antigen 1
EMT: epithelial- mesenchymal transition
ERK1/2: extracellular signal-regulated kinase 1/2
FBS: fetal bovine serum
FGM: full growth medium
GFP: green fluorescent protein
GO: gene ontology
HEK: human embryonic kidney
IFM: immunofluorescence microscopy
IFT: intraflagellar transport
ipTM: interface predicted template modeling score
KO: knockout
LC3B: microtubule- associated protein 1 light chain 3 beta
PAE: predicted aligned error
PCC: Pearson’s correlation coefficient
PDB: Protein Data Bank
qPCR: quantitative polymerase chain reaction
RAB7A: RAS-related protein RAB7A
RAB11B: RAS-related protein RAB11B
RIPA: radioimmunoprecipitation assay buffer
RPE: retinal pigment epithelium
SEM: standard error of the mean
SMAD2/3: small mothers against decapentaplegic homologues 2 and 3
SS: serum-starved medium
TGF-β: transforming growth factor beta
TGFBR1: transforming growth factor beta receptor type 1
WB: Western blot
WT: wild-type

## Declarations

### Ethics approval and consent to participate

Not applicable.

## Consent for publication

Not applicable.

## Availability of data and materials

The materials used and generated in this study (i.e. plasmids, cell lines, etc.) are available from the lead contact. Further information and requests for resources and additional data should be directed to and will be fulfilled by the lead contact, Diana Valverde. The mass spectrometry proteomics data have been deposited in the ProteomeXchange Consortium via PRIDE under dataset identifier PXD057025 (10.6019/PXD057025).

## Authors’ contributions

Conceptualization, C.S and D.V; methodology, C.S., J.B-C, S.V-A, C.A, P.B, and M.S; writing, all authors; funding acquisition, D.V, N.G and S.T.C; supervision, D.V, N.G. and S.T.C.

## Acknowledgements

We would like to thank Alexander von Kriegsheim and Roopesh Krishnankutty from the HTPU and Mass Spectrometry Facility of Cancer Research UK Edinburgh Centre for their help and efficiency in processing the samples in the mass spectrometer. Also, to Alina Zitskaja, Agata Makar, and Joanne Simpson from Noor Gammoh’s lab for their technical help. Thanks to Ainhoa Rodríguez Tébar, from the confocal microscopy service of the Galicia Sur Health Research Institute, for her help with obtaining images. To Mercedes Peleteiro Olmedo, from the cytometry service at CINBIO, for her help in handling the clones. CACTI Sequencing service Ángel Sebastián Comesaña and Verónica Outeiriño. CACTI Proteomic service: Paula Chaver and Manuel Marcos. Pedro Loureiro Pego for his help selecting clones. Mauro Lago Docampo for critically reviewing the manuscript. Ana Paula Borges Diez for her help designing the figures.

Authors’ information

## Supplementary data

**Supplementary Figure 1.**
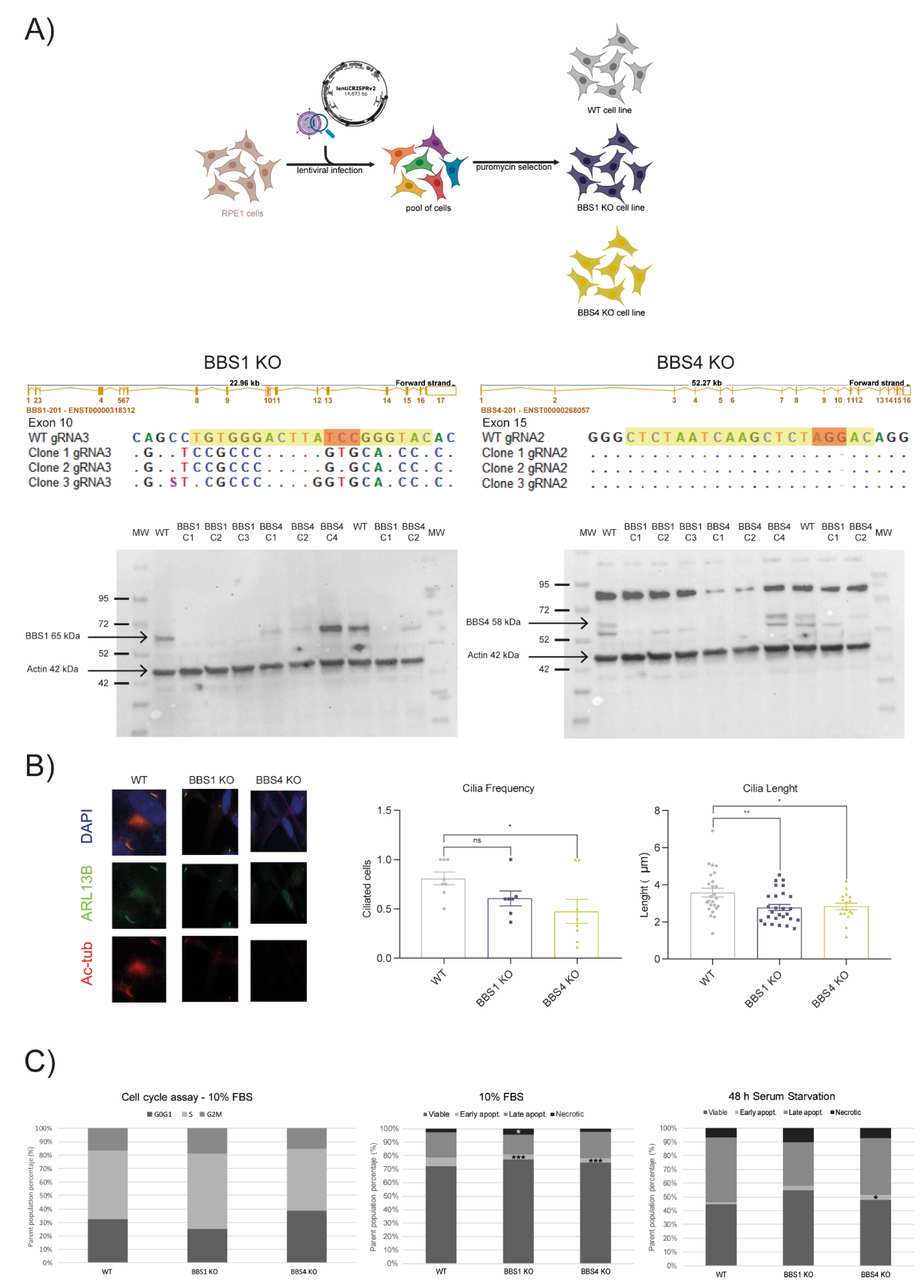
Validation and characterisation of BBS1 and BBS4 KO cell lines. (A) CRISPR-Cas9 targeting strategy: DNA sequences of KO clones for each gene, with the gRNA target region highlighted in yellow and the PAM sequence in red, and western blot validation of complete ablation of BBS1 and BBS4 proteins in the respective KO clones using β-actin as a loading control. (B) Representative immunofluorescence images of primary cilia stained with ARL13B and acetylated tubulin, and quantification of ciliary frequency and length in WT, BBS1 KO, and BBS4 KO cells following 24 h serum starvation. 4 fields were quantified per genotype across n = 3 independent biological replicates. (C) Cell cycle distribution assessed by flow cytometry under 10% FBS conditions, and apoptosis/viability assessed by Annexin V/propidium iodide staining under 10% FBS and 48 h serum starvation. Data are from n = 3 independent biological replicates. The differences were considered significant when *p < 0.05, **p < 0.01, and ***p < 0.001. ns = non-significant.

**Supplementary Figure 2.**
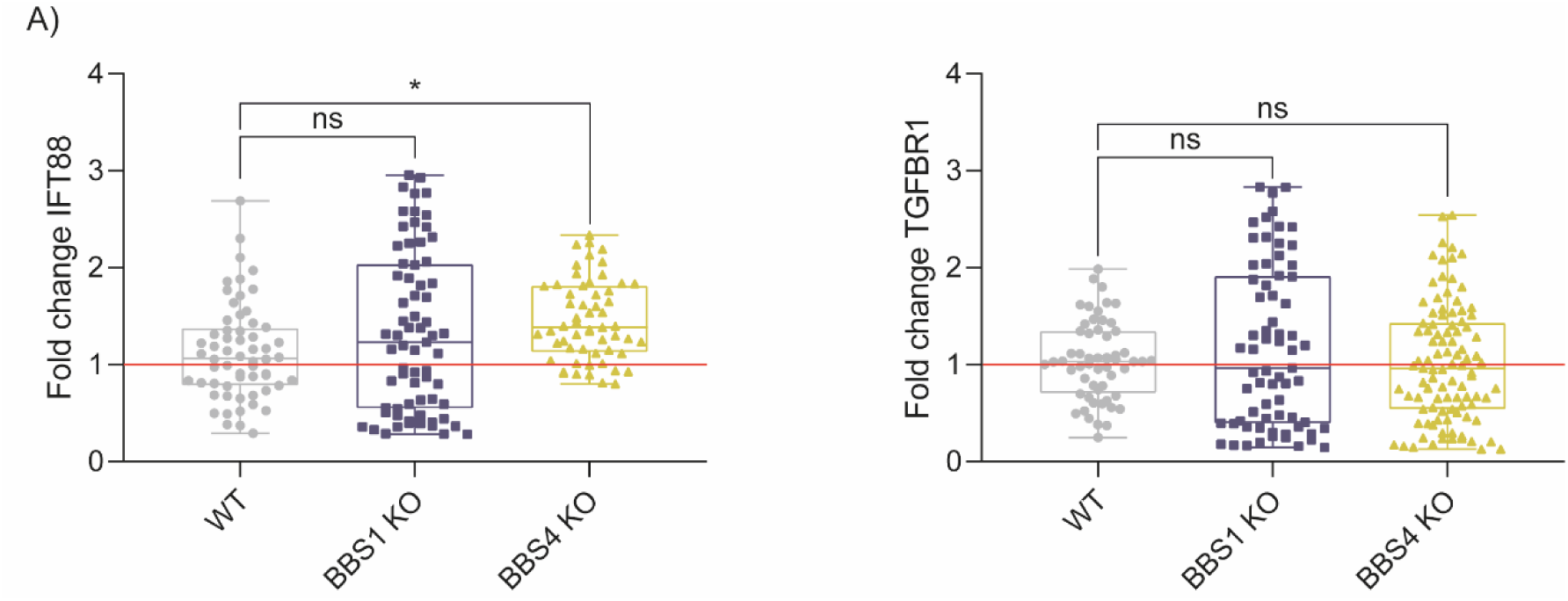
Validation of TGFBR1 and IFT88 gene expression in KO cell lines. qPCR analysis of TGFBR1 and IFT88 transcript levels in WT, BBS1 KO, and BBS4 KO cells, confirming that accumulation of TGFBR1 at the plasma membrane is not caused by transcriptional overexpression, and that the observed phenotypes are not attributable to disruption of general ciliary transport mechanisms. Each replicate was measured in technical triplicate and normalised to WT. Data represent mean ± SEM from n = 22 across n = 3 independent biological replicates. ns = non-significant.

## References

1. Satir P, Christensen ST. Overview of structure and function of mammalian cilia. Annu Rev Physiol. 2007;69:377–400. doi:10.1146/annurev.physiol.69.040705.141236 PubMed PMID: 17009929.

2. Anvarian Z, Mykytyn K, Mukhopadhyay S, Pedersen LB, Christensen ST. Cellular signalling by primary cilia in development, organ function and disease. Nat Rev Nephrol. 2019 Apr;15(4):199–219. doi:10.1038/s41581-019-0116-9 PubMed PMID: 30733609; PubMed Central PMCID: PMC6426138.

3. Hilgendorf KI, Myers BR, Reiter JF. Emerging mechanistic understanding of cilia function in cellular signalling. Nat Rev Mol Cell Biol. 2024 Jul;25(7):555–73. doi:10.1038/s41580-023-00698-5 PubMed PMID: 38366037; PubMed Central PMCID: PMC11199107.

4. Reiter JF, Leroux MR. Genes and molecular pathways underpinning ciliopathies. Nat Rev Mol Cell Biol. 2017 Sep;18(9):533–47. doi:10.1038/nrm.2017.60 PubMed PMID: 28698599; PubMed Central PMCID: PMC5851292.

5. Mill P, Christensen ST, Pedersen LB. Primary cilia as dynamic and diverse signalling hubs in development and disease. Nat Rev Genet. 2023 Jul;24(7):421–41. doi:10.1038/s41576-023-00587-9 PubMed PMID: 37072495; PubMed Central PMCID: PMC7615029.

6. Denniston AK, Beales PL, Tomlins PJ, Good P, Langford M, Foggensteiner L, et al. Evaluation of visual function and needs in adult patients with bardet-biedl syndrome. Retina Phila Pa. 2014 Nov;34(11):2282–9. doi:10.1097/IAE.0000000000000222 PubMed PMID: 25170860.

7. Forsythe E, Kenny J, Bacchelli C, Beales PL. Managing Bardet–Biedl Syndrome— Now and in the Future. Front Pediatr. 2018 Feb 13;6:23. doi:10.3389/fped.2018.00023 PubMed PMID: 29487844; PubMed Central PMCID: PMC5816783.

8. Wingfield JL, Lechtreck KF, Lorentzen E. Trafficking of ciliary membrane proteins by the intraflagellar transport/BBSome machinery. Essays Biochem. 2018 Dec 7;62(6):753– 63. doi:10.1042/EBC20180030 PubMed PMID: 30287585; PubMed Central PMCID: PMC6737936.

9. Álvarez-Satta M, Lago-Docampo M, Bea-Mascato B, Solarat C, Castro-Sánchez S, Christensen ST, et al. ALMS1 Regulates TGF-β Signaling and Morphology of Primary Cilia. Front Cell Dev Biol. 2021;9:623829. doi:10.3389/fcell.2021.623829 PubMed PMID: 33598462; PubMed Central PMCID: PMC7882606.

10. Westlake CJ, Baye LM, Nachury MV, Wright KJ, Ervin KE, Phu L, et al. Primary cilia membrane assembly is initiated by Rab11 and transport protein particle II (TRAPPII) complex-dependent trafficking of Rabin8 to the centrosome. Proc Natl Acad Sci U S A. 2011 Feb 15;108(7):2759–64. doi:10.1073/pnas.1018823108 PubMed PMID: 21273506; PubMed Central PMCID: PMC3041065.

11. Benmerah A. The ciliary pocket. Curr Opin Cell Biol. 2013 Feb;25(1):78–84. doi:10.1016/j.ceb.2012.10.011 PubMed PMID: 23153502.

12. Clement CA, Ajbro KD, Koefoed K, Vestergaard ML, Veland IR, Henriques de Jesus MPR, et al. TGF-β signaling is associated with endocytosis at the pocket region of the primary cilium. Cell Rep. 2013 Jun 27;3(6):1806–14. doi:10.1016/j.celrep.2013.05.020 PubMed PMID: 23746451.

13. Bhattacharyya S, Rainey MA, Arya P, Mohapatra BC, Mushtaq I, Dutta S, et al. Endocytic recycling protein EHD1 regulates primary cilia morphogenesis and SHH signaling during neural tube development. Sci Rep. 2016 Feb 17;6:20727. doi:10.1038/srep20727 PubMed PMID: 26884322; PubMed Central PMCID: PMC4756679.

14. Mönnich M, Borgeskov L, Breslin L, Jakobsen L, Rogowski M, Doganli C, et al. CEP128 Localizes to the Subdistal Appendages of the Mother Centriole and Regulates TGF-β/BMP Signaling at the Primary Cilium. Cell Rep. 2018 Mar 6;22(10):2584–92. doi:10.1016/j.celrep.2018.02.043 PubMed PMID: 29514088.

15. Barral DC, Garg S, Casalou C, Watts GFM, Sandoval JL, Ramalho JS, et al. Arl13b regulates endocytic recycling traffic. Proc Natl Acad Sci U S A. 2012 Dec 26;109(52):21354–9. doi:10.1073/pnas.1218272110 PubMed PMID: 23223633; PubMed Central PMCID: PMC3535586.

16. Seo S, Guo DF, Bugge K, Morgan DA, Rahmouni K, Sheffield VC. Requirement of Bardet-Biedl syndrome proteins for leptin receptor signaling. Hum Mol Genet. 2009 Apr 1;18(7):1323–31. doi:10.1093/hmg/ddp031 PubMed PMID: 19150989; PubMed Central PMCID: PMC2655773.

17. Leitch CC, Lodh S, Prieto-Echagüe V, Badano JL, Zaghloul NA. Basal body proteins regulate Notch signaling through endosomal trafficking. J Cell Sci. 2014 Jun 1;127(11):2407–19. doi:10.1242/jcs.130344 PubMed PMID: 24681783; PubMed Central PMCID: PMC4038940.

18. Starks RD, Beyer AM, Guo DF, Boland L, Zhang Q, Sheffield VC, et al. Regulation of Insulin Receptor Trafficking by Bardet Biedl Syndrome Proteins. PLoS Genet. 2015 Jun;11(6):e1005311. doi:10.1371/journal.pgen.1005311 PubMed PMID: 26103456; PubMed Central PMCID: PMC4478011.

19. Guo DF, Cui H, Zhang Q, Morgan DA, Thedens DR, Nishimura D, et al. The BBSome Controls Energy Homeostasis by Mediating the Transport of the Leptin Receptor to the Plasma Membrane. PLoS Genet. 2016 Feb;12(2):e1005890. doi:10.1371/journal.pgen.1005890 PubMed PMID: 26926121; PubMed Central PMCID: PMC4771807.

20. Strauss O. The retinal pigment epithelium in visual function. Physiol Rev. 2005 Jul;85(3):845–81. doi:10.1152/physrev.00021.2004 PubMed PMID: 15987797.

21. Zhou M, Geathers JS, Grillo SL, Weber SR, Wang W, Zhao Y, et al. Role of Epithelial- Mesenchymal Transition in Retinal Pigment Epithelium Dysfunction. Front Cell Dev Biol. 2020 Jun 25;8:501. doi:10.3389/fcell.2020.00501 PubMed PMID: 32671066; PubMed Central PMCID: PMC7329994.

22. Ran FA, Hsu PD, Wright J, Agarwala V, Scott DA, Zhang F. Genome engineering using the CRISPR-Cas9 system. Nat Protoc. 2013 Nov;8(11):2281–308. doi:10.1038/nprot.2013.143 PubMed PMID: 24157548; PubMed Central PMCID: PMC3969860.

23. Shalem O, Sanjana NE, Hartenian E, Shi X, Scott DA, Mikkelson T, et al. Genome- scale CRISPR-Cas9 knockout screening in human cells. Science. 2014 Jan 3;343(6166):84–7. doi:10.1126/science.1247005 PubMed PMID: 24336571; PubMed Central PMCID: PMC4089965.

24. Yu G, Wang LG, Han Y, He QY. clusterProfiler: an R package for comparing biological themes among gene clusters. Omics J Integr Biol. 2012 May;16(5):284–7. doi:10.1089/omi.2011.0118 PubMed PMID: 22455463; PubMed Central PMCID: PMC3339379.

25. Walter W, Sánchez-Cabo F, Ricote M. GOplot: an R package for visually combining expression data with functional analysis. Bioinforma Oxf Engl. 2015 Sep 1;31(17):2912– 4. doi:10.1093/bioinformatics/btv300 PubMed PMID: 25964631.

26. J S, I AC, E F, V K, M L, T P, et al. Fiji: an open-source platform for biological-image analysis. Nat Methods. 2012 Jun 28;9(7). doi:10.1038/nmeth.2019 PubMed PMID: 22743772.

27. Prasai A, Schmidt Cernohorska M, Ruppova K, Niederlova V, Andelova M, Draber P, et al. The BBSome assembly is spatially controlled by BBS1 and BBS4 in human cells. J Biol Chem. 2020 Oct 16;295(42):14279–90. doi:10.1074/jbc.RA120.013905 PubMed PMID: 32759308; PubMed Central PMCID: PMC7573277.

28. Castro-Sánchez S, Suarez-Bregua P, Novas R, Álvarez-Satta M, Badano JL, Rotllant J, et al. Functional analysis of new human Bardet-Biedl syndrome loci specific variants in the zebrafish model. Sci Rep. 2019 Sep 10;9(1):12936. doi:10.1038/s41598-019-49217-7 PubMed PMID: 31506453; PubMed Central PMCID: PMC6736949.

29. Xie C, Habif JC, Uytingco CR, Ukhanov K, Zhang L, de Celis C, et al. Gene therapy rescues olfactory perception in a clinically relevant ciliopathy model of Bardet-Biedl syndrome. FASEB J Off Publ Fed Am Soc Exp Biol. 2021 Sep;35(9):e21766. doi:10.1096/fj.202100627R PubMed PMID: 34383976; PubMed Central PMCID: PMC8785972.

30. Castro-Sánchez S, Álvarez-Satta M, Cortón M, Guillén E, Ayuso C, Valverde D. Exploring genotype-phenotype relationships in Bardet-Biedl syndrome families. J Med Genet. 2015 Aug;52(8):503–13. doi:10.1136/jmedgenet-2015-103099 PubMed PMID: 26082521.

31. Niederlova V, Modrak M, Tsyklauri O, Huranova M, Stepanek O. Meta-analysis of genotype-phenotype associations in Bardet-Biedl syndrome uncovers differences among causative genes. Hum Mutat. 2019 Nov;40(11):2068–87. doi:10.1002/humu.23862 PubMed PMID: 31283077.

32. Bökel C, Schwabedissen A, Entchev E, Renaud O, González-Gaitán M. Sara endosomes and the maintenance of Dpp signaling levels across mitosis. Science. 2006 Nov 17;314(5802):1135–9. doi:10.1126/science.1132524 PubMed PMID: 17110576.

33. Coumailleau F, Fürthauer M, Knoblich JA, González-Gaitán M. Directional Delta and Notch trafficking in Sara endosomes during asymmetric cell division. Nature. 2009 Apr 23;458(7241):1051–5. doi:10.1038/nature07854 PubMed PMID: 19295516.

34. Devenport D, Oristian D, Heller E, Fuchs E. Mitotic internalization of planar cell polarity proteins preserves tissue polarity. Nat Cell Biol. 2011 Jul 10;13(8):893–902. doi:10.1038/ncb2284 PubMed PMID: 21743464; PubMed Central PMCID: PMC3149741.

35. Cota CD, Davidson B. Mitotic Membrane Turnover Coordinates Differential Induction of the Heart Progenitor Lineage. Dev Cell. 2015 Sep 14;34(5):505–19. doi:10.1016/j.devcel.2015.07.001 PubMed PMID: 26300448.

36. Heck BW, Devenport D. Trans-endocytosis of Planar Cell Polarity Complexes during Cell Division. Curr Biol CB. 2017 Dec 4;27(23):3725–3733.e4. doi:10.1016/j.cub.2017.10.053 PubMed PMID: 29174888; PubMed Central PMCID: PMC5728440.

37. D J, G C, J K, E T, G N, C LG, et al. Alström Syndrome protein ALMS1 localizes to basal bodies of cochlear hair cells and regulates cilium-dependent planar cell polarity. Hum Mol Genet. 2011 Jan 2;20(3). doi:10.1093/hmg/ddq493 PubMed PMID: 21071598.

38. Hl MS, Jd G, C G, M C, Sm C, T B, et al. Loss of MACF1 Abolishes Ciliogenesis and Disrupts Apicobasal Polarity Establishment in the Retina. Cell Rep. 2016 Oct 25;17(5). doi:10.1016/j.celrep.2016.09.089 PubMed PMID: 27783952.

39. Wang L, Liu Y, Stratigopoulos G, Panigrahi S, Sui L, Zhang Y, et al. Bardet-Biedl syndrome proteins regulate intracellular signaling and neuronal function in patient-specific iPSC-derived neurons. J Clin Invest. 2021 Apr 15;131(8):146287. doi:10.1172/JCI146287 PubMed PMID: 33630762; PubMed Central PMCID: PMC8262481.

40. Taschner M, Lorentzen E. The Intraflagellar Transport Machinery. Cold Spring Harb Perspect Biol. 2016 Oct 3;8(10):a028092. doi:10.1101/cshperspect.a028092 PubMed PMID: 27352625; PubMed Central PMCID: PMC5046692.

41. Lin YH, Currinn H, Pocha SM, Rothnie A, Wassmer T, Knust E. AP-2-complex- mediated endocytosis of Drosophila Crumbs regulates polarity by antagonizing Stardust. J Cell Sci. 2015 Dec 15;128(24):4538–49. doi:10.1242/jcs.174573 PubMed PMID: 26527400.

42. Goto H, Inaba H, Inagaki M. Mechanisms of ciliogenesis suppression in dividing cells. Cell Mol Life Sci CMLS. 2017 Mar;74(5):881–90. doi:10.1007/s00018-016-2369-9 PubMed PMID: 27669693; PubMed Central PMCID: PMC5306231.

43. Lönn P, Morén A, Raja E, Dahl M, Moustakas A. Regulating the stability of TGFβ receptors and Smads. Cell Res. 2009 Jan;19(1):21–35. doi:10.1038/cr.2008.308

44. Huang F, Chen YG. Regulation of TGF-β receptor activity. Cell Biosci. 2012 Mar 15;2(1):9. doi:10.1186/2045-3701-2-9

45. Zhu L, Chen S, Chen Y. Unraveling the biological functions of Smad7 with mouse models. Cell Biosci. 2011 Dec 28;1(1):44. doi:10.1186/2045-3701-1-44

46. S G, N O, J K, S PS, R DP, M D, et al. TGF-β receptor I/II trafficking and signaling at primary cilia are inhibited by ceramide to attenuate cell migration and tumor metastasis. Sci Signal. 2017 Oct 24;10(502). doi:10.1126/scisignal.aam7464 PubMed PMID: 29066540.

47. Kavsak P, Rasmussen RK, Causing CG, Bonni S, Zhu H, Thomsen GH, et al. Smad7 Binds to Smurf2 to Form an E3 Ubiquitin Ligase that Targets the TGFβ Receptor for Degradation. Mol Cell. 2000 Dec 1;6(6):1365–75. doi:10.1016/S1097-2765(00)00134-9

48. O’Brien CE, Bonanno L, Zhang H, Wyss-Coray T. Beclin 1 regulates neuronal transforming growth factor-β signaling by mediating recycling of the type I receptor ALK5. Mol Neurodegener. 2015 Dec 21;10:69. doi:10.1186/s13024-015-0065-0 PubMed PMID: 26692002.

49. Ishikawa K, Sreekumar PG, Spee C, Nazari H, Zhu D, Kannan R, et al. αB-Crystallin Regulates Subretinal Fibrosis by Modulation of Epithelial-Mesenchymal Transition. Am J Pathol. 2016 Apr;186(4):859–73. doi:10.1016/j.ajpath.2015.11.014 PubMed PMID: 26878210; PubMed Central PMCID: PMC4822331.

50. Schneider S, De Cegli R, Nagarajan J, Kretschmer V, Matthiessen PA, Intartaglia D, et al. Loss of Ciliary Gene Bbs8 Results in Physiological Defects in the Retinal Pigment Epithelium. Front Cell Dev Biol. 2021;9:607121. doi:10.3389/fcell.2021.607121 PubMed PMID: 33681195; PubMed Central PMCID: PMC7930748.

51. Weihbrecht K, Goar WA, Pak T, Garrison JE, DeLuca AP, Stone EM, et al. Keeping an Eye on Bardet-Biedl Syndrome: A Comprehensive Review of the Role of Bardet-Biedl Syndrome Genes in the Eye. Med Res Arch. 2017 Sep;5(9). doi:10.18103/mra.v5i9.1526 PubMed PMID: 29457131; PubMed Central PMCID: PMC5814251.

52. Hsu Y, Seo S, Sheffield VC. Photoreceptor cilia, in contrast to primary cilia, grant entry to a partially assembled BBSome. Hum Mol Genet. 2021 Mar 25;30(1):87–102. doi:10.1093/hmg/ddaa284 PubMed PMID: 33517424; PubMed Central PMCID: PMC8033148.

53. Langousis G, Shimogawa MM, Saada EA, Vashisht AA, Spreafico R, Nager AR, et al. Loss of the BBSome perturbs endocytic trafficking and disrupts virulence of Trypanosoma brucei. Proc Natl Acad Sci U S A. 2016 Jan 19;113(3):632–7. doi:10.1073/pnas.1518079113 PubMed PMID: 26721397; PubMed Central PMCID: PMC4725476.

54. Zhang X, Liu J, Pan T, Ward A, Liu J, Xu XZS. A cilia-independent function of BBSome mediated by DLK-MAPK signaling in C. elegans photosensation. Dev Cell. 2022 Jun 20;57(12):1545–1557.e4. doi:10.1016/j.devcel.2022.05.005 PubMed PMID: 35649417; PubMed Central PMCID: PMC9233093.

55. Perron M, Boy S, Amato MA, Viczian A, Koebernick K, Pieler T, et al. A novel function for Hedgehog signalling in retinal pigment epithelium differentiation. Dev Camb Engl. 2003 Apr;130(8):1565–77. doi:10.1242/dev.00391 PubMed PMID: 12620982.

56. Burke JM. Epithelial phenotype and the RPE: is the answer blowing in the Wnt? Prog Retin Eye Res. 2008 Nov;27(6):579–95. doi:10.1016/j.preteyeres.2008.08.002 PubMed PMID: 18775790; PubMed Central PMCID: PMC2584165.

57. Schouwey K, Aydin IT, Radtke F, Beermann F. RBP-Jκ-dependent Notch signaling enhances retinal pigment epithelial cell proliferation in transgenic mice. Oncogene. 2011 Jan 20;30(3):313–22. doi:10.1038/onc.2010.428 PubMed PMID: 20856205.

58. Haws RM, Gordon G, Han JC, Yanovski JA, Yuan G, Stewart MW. The efficacy and safety of setmelanotide in individuals with Bardet-Biedl syndrome or Alström syndrome: Phase 3 trial design. Contemp Clin Trials Commun. 2021 Jun;22:100780. doi:10.1016/j.conctc.2021.100780 PubMed PMID: 34013094; PubMed Central PMCID: PMC8114053.

59. Koo BK, Spit M, Jordens I, Low TY, Stange DE, van de Wetering M, et al. Tumour suppressor RNF43 is a stem-cell E3 ligase that induces endocytosis of Wnt receptors. Nature. 2012 Aug 30;488(7413):665–9. doi:10.1038/nature11308 PubMed PMID: 22895187.

60. Jin H, White SR, Shida T, Schulz S, Aguiar M, Gygi SP, et al. The conserved Bardet- Biedl syndrome proteins assemble a coat that traffics membrane proteins to cilia. Cell. 2010 Jun 25;141(7):1208–19. doi:10.1016/j.cell.2010.05.015 PubMed PMID: 20603001; PubMed Central PMCID: PMC2898735.

61. Klink BU, Gatsogiannis C, Hofnagel O, Wittinghofer A, Raunser S. Structure of the human BBSome core complex. eLife. 2020 Jan 17;9:e53910. doi:10.7554/eLife.53910 PubMed PMID: 31951201; PubMed Central PMCID: PMC7018512.

62. Tian X, Zhao H, Zhou J. Organization, functions, and mechanisms of the BBSome in development, ciliopathies, and beyond. eLife. 2023 Jul 19;12:e87623. doi:10.7554/eLife.87623 PubMed PMID: 37466224.

63. Di Guglielmo GM, Le Roy C, Goodfellow AF, Wrana JL. Distinct endocytic pathways regulate TGF-beta receptor signalling and turnover. Nat Cell Biol. 2003 May;5(5):410–21. doi:10.1038/ncb975 PubMed PMID: 12717440.

64. Chen YG. Endocytic regulation of TGF-β signaling. Cell Res. 2009 Jan;19(1):58–70. doi:10.1038/cr.2008.315

65. He K, Yan X, Li N, Dang S, Xu L, Zhao B, et al. Internalization of the TGF-β type I receptor into caveolin-1 and EEA1 double-positive early endosomes. Cell Res. 2015 Jun;25(6):738–52. doi:10.1038/cr.2015.60

66. Balogh P, Katz S, Kiss AL. The role of endocytic pathways in TGF-β signaling. Pathol Oncol Res POR. 2013 Apr;19(2):141–8. doi:10.1007/s12253-012-9595-8 PubMed PMID: 23274761.

67. Ehrlich M. Endocytosis and trafficking of BMP receptors: Regulatory mechanisms for fine-tuning the signaling response in different cellular contexts. Cytokine Growth Factor Rev. 2016 Feb;27:35–42. doi:10.1016/j.cytogfr.2015.12.008 PubMed PMID: 26776724.

68. Pedersen LB, Mogensen JB, Christensen ST. Endocytic Control of Cellular Signaling at the Primary Cilium. Trends Biochem Sci. 2016 Sep;41(9):784–97. doi:10.1016/j.tibs.2016.06.002 PubMed PMID: 27364476.

69. Vander Ark A, Cao J, Li X. TGF-β receptors: In and beyond TGF-β signaling. Cell Signal. 2018 Dec;52:112–20. doi:10.1016/j.cellsig.2018.09.002 PubMed PMID: 30184463.

70. Prieto-Echagüe V, Lodh S, Colman L, Bobba N, Santos L, Katsanis N, et al. BBS4 regulates the expression and secretion of FSTL1, a protein that participates in ciliogenesis and the differentiation of 3T3-L1. Sci Rep. 2017 Aug 29;7(1):9765. doi:10.1038/s41598-017-10330-0 PubMed PMID: 28852127; PubMed Central PMCID: PMC5575278.

71. Derivery E, Seum C, Daeden A, Loubéry S, Holtzer L, Jülicher F, et al. Polarized endosome dynamics by spindle asymmetry during asymmetric cell division. Nature. 2015 Dec 10;528(7581):280–5. doi:10.1038/nature16443 PubMed PMID: 26659188.

72. Schmid FM, Schou KB, Vilhelm MJ, Holm MS, Breslin L, Farinelli P, et al. IFT20 modulates ciliary PDGFRα signaling by regulating the stability of Cbl E3 ubiquitin ligases. J Cell Biol. 2018 Jan 2;217(1):151–61. doi:10.1083/jcb.201611050 PubMed PMID: 29237719; PubMed Central PMCID: PMC5748969.

73. Guo DF, Rahmouni K. The Bardet-Biedl syndrome protein complex regulates cell migration and tissue repair through a Cullin-3/RhoA pathway. Am J Physiol Cell Physiol. 2019 Sep 1;317(3):C457–65. doi:10.1152/ajpcell.00498.2018 PubMed PMID: 31216194; PubMed Central PMCID: PMC6766617.

74. Freke GM, Martins T, Davies RJ, Beyer T, Seda M, Peskett E, et al. De-Suppression of Mesenchymal Cell Identities and Variable Phenotypic Outcomes Associated with Knockout of Bbs1. Cells. 2023 Nov 20;12(22):2662. doi:10.3390/cells12222662 PubMed PMID: 37998397; PubMed Central PMCID: PMC10670506.

75. Kim SJ, Kim YS, Kim JH, Jang HY, Ly DD, Das R, et al. Activation of ERK1/2- mTORC1-NOX4 mediates TGF-β1-induced epithelial-mesenchymal transition and fibrosis in retinal pigment epithelial cells. Biochem Biophys Res Commun. 2020 Aug 27;529(3):747–52. doi:10.1016/j.bbrc.2020.06.034 PubMed PMID: 32736702.

76. Huang Y, Meister R, Lindziute M, Binter M, Tode J, Framme C, et al. A TGFB2/TNF- induced in vitro model of proliferative vitreoretinopathy (PVR) using ARPE-19 cells confirms nicotinamide as an inhibitor of EMT and VEGFA secretion. PloS One. 2026;21(1):e0340614. doi:10.1371/journal.pone.0340614 PubMed PMID: 41529022; PubMed Central PMCID: PMC12798965.

