## Supplementary material for "Depletion of BBSome Subunits Alters Receptor Endocytosis and Promotes EMT via TGF-β Signalling": Supl_material: Supplementary Table 1 , 2 and 3.docx

**Supplementary Table S1.** sgRNAs used to generate knockout cell lines in RPE1 for *BBS1* and *BBS4* genes.

| **ID** | **Gene** | **Exon** | **Forward primer (5' - 3')** | **Reverse primer (5' - 3')** | **Strand** | **PAM** | **On-target Score** | **Off-target Score** |
| --- | --- | --- | --- | --- | --- | --- | --- | --- |
| gRNA 1 | *BBS1* | 9 | CACCGTGTTGAGTTCCGGCTTGCCG | AAACCGGCAAGCCGGAACTCAACAC | + | CGG | 61,7 | 92 |
| gRNA 2 | *BBS1* | 1 | CACCGCAGGCGTCGGAATCCGATG | AAACCATCGGATTCCGACGCCTGC | - | AGG | 69,2 | 92,4 |
| gRNA 3 | *BBS1* | 10 | CACCGTGTACCCGGATAAGTCCCAC | AAACGTGGGACTTATCCGGGTACAC | - | AGG | 65 | 83 |
| gRNA 4 | *BBS1* | 4 | CACCGGGCTTTCGGTCATCACCAG | AAACCTGGTGATGACCGAAAGCCC | - | TGG | 71,3 | 75,3 |
| gRNA 1 | *BBS4* | 1 | CACCGAACGCCCTGGCCCAATATC | AAACGATATTGGGCCAGGGCGTTC | + | GGG | 42,9 | 90,5 |
| gRNA 2 | *BBS4* | 7 | CACCGCTCTAATCAAGCTCTAGGAC | AAACGTCCTAGAGCTTGATTAGAGC | - | TGG | 44,2 | 78,9 |
| gRNA 3 | *BBS4* | Intron 1-2 | CACCGTGTCTTCAGCAGCTGCATAC | AAACGTATGCAGCTGCTGAAGACAC | + | GGG | 50,1 | 95,6 |
| gRNA 4 | *BBS4* | Intron 1-2 | CACCGTGAGGTTTATTAGTCCCCA | AAACTGGGGACTAATAAACCTCAC | + | GGG | 50,4 | 94,0 |
| V6 |  |  | GACTATCATATGCTTACCGT |  |  |  |  |  |

**Supplementary Table S2.** Primers used to check the effects of the CRISPR-Cas9 process in the knockout cell lines.

| Primer name | Sequence | Tm (ºC) |
| --- | --- | --- |
| BBS1_gRNA1_FWD | TGGGACTTTAGACCAGGCAC | 57 |
| BBS1_gRNA1_RVS | AAAGCCCACTCTCATCTTGC | 57 |
| BBS1_gRNA2_FWD | TGAGCCTGGGTGGGAAAG | 57 |
| BBS1_gRNA2_RVS | AGGCCTCGGTTTCCCTATCT | 57 |
| BBS1_gRNA3_FWD | AAGGTGGCAGAAGTGGAAAT | 57 |
| BBS1_gRNA3_RVS | AGGGAACAGCCAGCGTCT | 57 |
| BBS1_gRNA4_FWD | AAGCCTTTATTGGTGCAGGA | 57 |
| BBS1_gRNA4_RVS | TAGACACAAGGGCCTGAAGC | 57 |
| BBS1_qPCR_FWD | TCCCCGTCTTCCTAGAGGTT | 60 |
| BBS1_qPCR_RVS | TCGATGCAGTACTTGGGGTG | 60 |
| BBS4_gRNA1_FWD | TCTGTCTGCCACAGGTGTAAC | 57 |
| BBS4_gRNA1_RVS | CCAGAACAGCCATGAACGTG | 57 |
| BBS4_gRNA2+3_FWD | GGTGGAGATGGCTCAGAAGT | 57 |
| BBS4_gRNA2+3_RVS | TCTCCCCTTGTGGCCAATAC | 57 |
| BBS4_gRNA4_FWD | ACAGACCTCTGCTGAGTAGC | 57 |
| BBS4_gRNA4_RVS | GCAATACTCAGCAGGCTTGG | 57 |
| BBS4_qPCR_FWD | TCAAGCAGGTGGCCAGATCT | 60 |
| BBS4_qPCR_RVS | GGTTATGGCTGATCTCCCAATC | 60 |

**Supplementary Table S3.** Primers used to study the expression of EMT and autophagy markers and to quantify the expression of the target genes in the knockout cell lines.

| Gene symbol | Forward primer | Reverse primer |
| --- | --- | --- |
| *CDH1* | GCCTCCTGAAAAGAGAGTGGAAG | TGGCAGTGTCTCTCCAAATCCG |
| *CDH2* | CCTCCAGAGTTTACTGCCATGAC | GTAGGATCTCCGCCACTGATTC |
| *ACTA2* | CTATGCCTCTGGACGCACAACT | CAGATCCAGACGCATGATGGCA |
| *SNAI1* | GCTGCAGGACTCTAATCCAGAGTT | GACAGAGTCCCAGATGAGCATTG |
| *SNAI2* | ATCTGCGGCAAGGCGTTTTCCA | GAGCCCTCAGATTTGACCTGTC |
| *VIM* | AGGCAAAGCAGGAGTCCACTGA | ATCTGGCGTTCCAGGGACTCAT |
| *VCL* | GAGCAAGCACAGCGGTGGATT | TCGGTCACACTTGGCGAGAAGA |
| *CRYAB* | ACTTCCCTGAGTCCCTTCTACC | GGAGAAGTGCTTCACATCCAGG |
| *CTNNB1* | CACAAGCAGAGTGCTGAAGGTG | GATTCCTGAGAGTCCAAAGACAG |
| *TWIST1* | GCCAGGTACATCGACTTCCTCT | TCCATCCTCCAGACCGAGAAGG |
| *IFT88* | TCGGCTAGATGAGGCTTTGGAC | CACTGACCACCTGCATTAGCCA |
| *TGFBR1* | GACAACGTCAGGTTCTGGCTCA | CCGCCACTTTCCTCTCCAAACT |
| *ULK1* | GCAAGGACTCTTCTTGTGACAC | CCACTGCACATCAGGCTGTCTG |
| *ATG5* | GCAGATGGACAGTTGCACACAC | GAGGTGTTTCCAACATTGGCTCA |
| *ATG7* | CGTTGCCCACAGCATCATCTTC | CACTGAGGTTCACCATCCTTGG |
| *BBS1* | TCCCCGTCTTCCTAGAGGTT | TCGATGCAGTACTTGGGGTG |
| *BBS4* | TCAAGCAGGTGGCCAGATCT | GGTTATGGCTGATCTCCCAATC |
| *ALAS1* | AGTGTGAAAACCGATGGAGG | CGATCATACTGAAAAGTGGAAACAG |
| *YWHAZ* | ATGCAACCAACACATCCTATC | GCATTATTAGCGTGCTGTCTT |
| *B2M* | TTTCATCCATCCGACATTGA | CCTCCATGATGCTGCTTACA |
