## Supplementary figures and images for "Depletion of BBSome Subunits Alters Receptor Endocytosis and Promotes EMT via TGF-β Signalling"

### Suppl_fig1_CRISPR_charact.tif

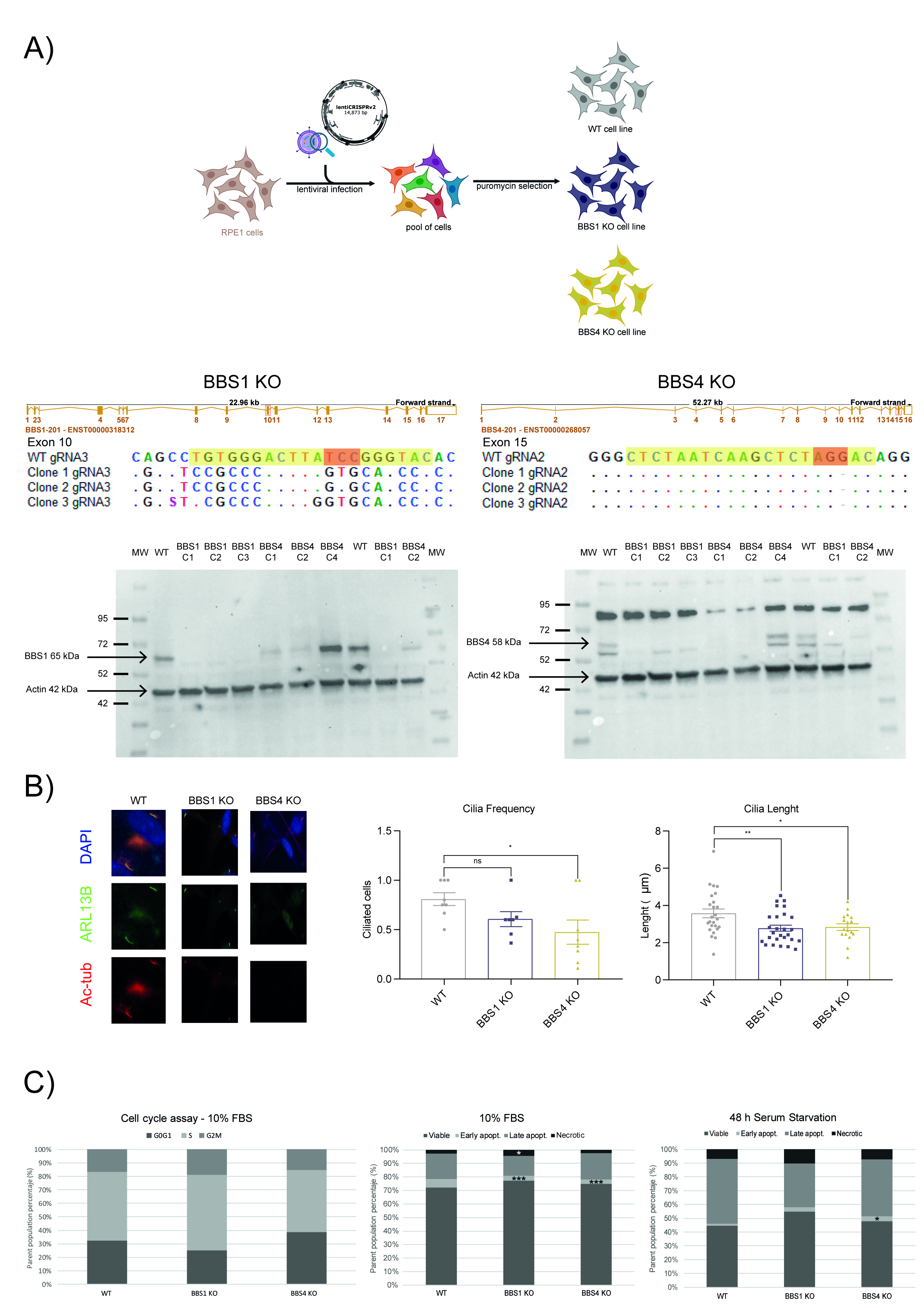

### Suppl_fig2.tif

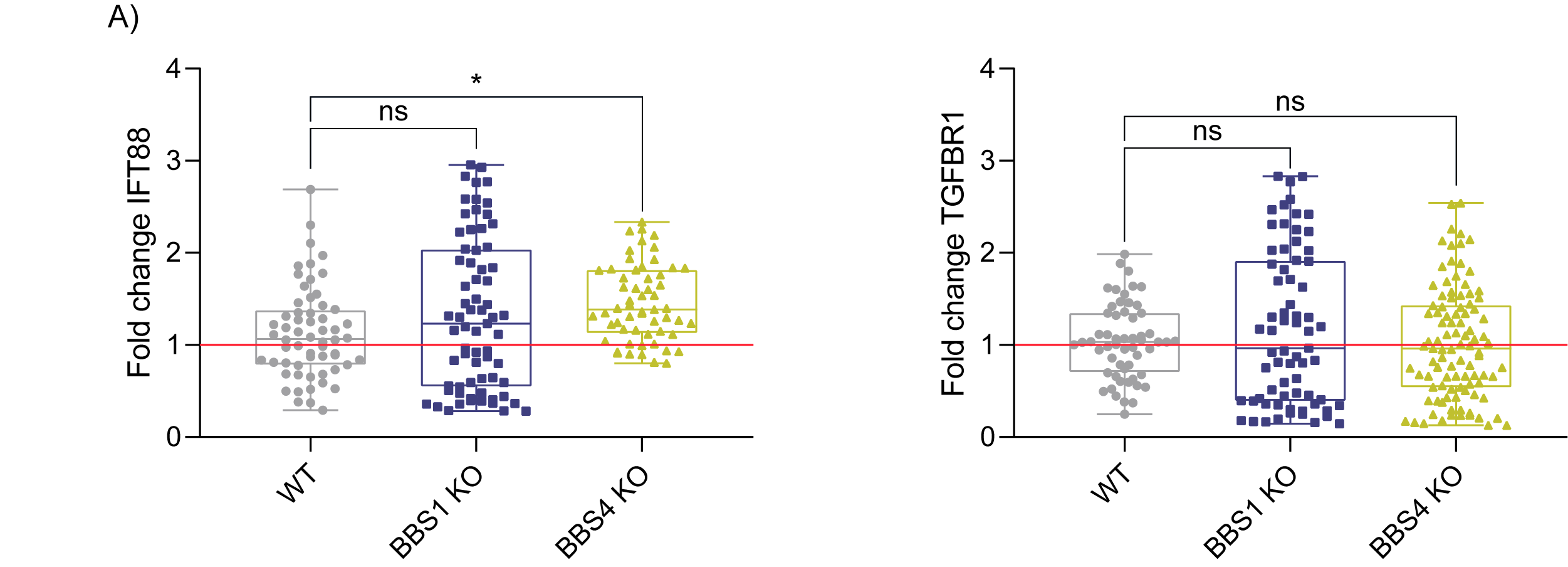
